# From vehicles to wildlife: transferable deep learning for trajectory generation

**DOI:** 10.64898/2026.07.14.736596

**Authors:** Julien Patras, Ronan Fablet, Adrien Brunel, Amédée Roy, Leandro Bugoni, Guilherme Tavares Nunes, Christophe Barbraud, Júlia Jacoby, Sophie Benboudjema, Giannina Passuni, Karine Delord, Sophie Lanco

## Abstract

Realistic simulation of animal movement is fundamental to conservation, habitat modeling, and ecological scenario evaluation. Traditional approaches struggle to capture multi-scale trajectory dynamics, while generative deep learning for complete trajectory simulation remains largely unexplored in ecology due to data scarcity. We show that a diffusion model pre-trained on millions of vehicle GPS trajectories can be fine-tuned on hundreds of seabird central-place foraging trips to simultaneously generate ecologically realistic trajectories for five species and six breeding colonies. The fine-tuned model consistently outperforms four state-of-the-art baselines (GAN, VAE, HMM, iSSF) across movement metrics spanning step dynamics, spatial distribution, and behavioral temporality, with comparable or shorter computation times. Domain transfer reaches full performance in 35 minutes versus 7 hours from scratch, and conditioning on species and colony enables generalization to unseen combinations from as few as 10 trajectories. These results establish cross-domain transfer learning as a new paradigm for data-efficient generative animal movement modeling.

## 1 Introduction

Every day, animals make movement decisions that determine their survival, their reproductive success, and ultimately the persistence of their populations. Predicting these decisions under accelerating environmental change is one of the central challenges of modern conservation [1], [2]. Recent advances in animal-borne telemetry have enabled the collection of high-frequency tracking data with increasingly negligible impacts on natural behavior, offering unprecedented insights into spatial ecology [3], [4], [5]. This massive data expansion has fostered the development of a wide range of modeling approaches aimed at understanding, describing, and simulating complex movement processes [2], [6], [7]. Movement models are central to simulating animal behavior, forecasting distribution shifts, and mitigating human-wildlife conflict under global change [1], [8], [9], [10].

Traditional movement modeling works predominantly from the bottom up, assuming that large-scale spatial patterns emerge from sequential, step-by-step transitions. While this framework has produced powerful tools for behavioral inference and habitat selection, it imposes a structural ceiling on realism: the trajectory as a whole is never modeled, only approximated as the sum of its steps. Correlated random walks and Lévy flights capture local statistical properties of movement but generally fail to reproduce coherent large-scale structure [11], [12]. Hidden Markov Models (HMMs) formalize behavioral state-switching at short timescales [13], [14], and integrated step selection functions (iSSFs) incorporate habitat covariates [15], [16], [9], [8]. Both, however, operate at the scale of individual steps, and neither reproduces large-scale movement structure. These approaches rely on the assumption that the underlying process is Markovian: the current state carries all the information needed to predict the next one, with no memory of what came before. Movement is a cohesive sequence of local decisions and energetic trade-offs that only becomes visible at the trajectory scale [17]: an animal may travel long distances in a coherent direction while foraging, then return to a central place once its energetic needs are met. This multi-scale structure matters because realistic simulation of complete trajectories underpins applications like predicting population redistribution under environmental change or assessing the spatial footprint of human activity. Even recent advances in habitat representation tend to produce trajectories with limited global realism [18]. Conversely, approaches that enforce large-scale constraints, such as home-range or central-place attraction, can reproduce some global structure, but usually at the cost of mechanistic detail and ecological realism [14], [19].

This limitation is particularly critical in systems where movement exhibits strong spatial and temporal organization across scales. In this context, seabirds provide a compelling case study: during the breeding season, they perform repeated round trips between their colony and distant foraging areas, known as central-place foraging trips (CPFTs). The structure of these trips emerges from the interplay of individual decision-making, prey distribution, oceanographic conditions, and energetic constraints [17], [20], [21]. Beyond their ecological interest, seabirds are key indicators of marine ecosystem dynamics due to their position at high trophic levels and their accessibility for observation and tracking [22], [23], [24]. At the same time, they are highly threatened, with a large share of species in decline due to multiple anthropogenic pressures [25]. Simulating their movement accurately is therefore essential for predicting their response to environmental change, evaluating conservation strategies such as marine protected areas, and assessing the impact of human activities like offshore renewable energy development. Current approaches to CPFT simulation rely mainly on HMMs [14], and more recently GANs [19], which outperform HMMs at large spatial scales but remain less accurate locally.

To capture these complex, trajectory-scale dynamics, deep learning frameworks have introduced a shifting paradigm in trajectory generation across other fields [26]. In autonomous vehicles and maritime navigation, models trained on millions of GPS tracks now produce diverse, spatially realistic, and physically plausible trajectories [27], [28]. Unlike step-by-step approaches, these methods generate the entire trajectory at once, treating it as a single object that can capture long-term dependencies. Among them, probabilistic diffusion denoising models (DDPMs) have emerged as a promising approach for generating complete trajectories that represent the state of the art in terms of diversity and fidelity [26].

The transformative impact of deep learning across many domains, such as industry and healthcare, has been driven by the rapid growth of available data and by increased interdisciplinarity between these fields and the communities of mathematicians and computer scientists. Ecology has likewise become a data-intensive discipline. Deep learning methods have therefore begun to spread within this field as well, although their applications are still largely focused on computer vision tasks [29], [30]. Topics such as animal behavior, movement, and species interactions have so far received comparatively less attention from the deep learning community [31]. However, training these architectures typically requires massive datasets, escalating as both model size and behavioral complexity increase. Recommended dataset sizes are typically not on the order of hundreds of samples, but rather tens of thousands or even millions of observations [30], [32]. In the life sciences, and more specifically in movement ecology, data collection is constrained by ethical, financial, and technical considerations, although the latter barrier is progressively being alleviated [33]. Species- and colony-specific datasets typically contain tens or hundreds of tracks, insufficient to train modern deep learning models from scratch. Nevertheless, it is possible to mitigate these constraints when applying data-driven model classes to small datasets by relying on techniques such as transfer learning, and more specifically, fine-tuning [34], [35]. By initializing from a model that has already incorporated general representations of spatiotemporal dynamics and movement structure, fine-tuning requires only a fraction of the data that would be needed for learning from scratch. While this approach has driven significant progress in natural language processing and computer vision [36], [37], [30], it has yet to be applied to animal movement simulation.

For nearly two decades, the movement ecology paradigm formulated by Nathan et al. [17] has provided the foundational framework for understanding the drivers and mechanisms of organismal movement. This paradigm was established at a time when animal movement was primarily studied through statistical analysis and hypothesis testing. While informative for specific features of the movement and explicit hypothesis testing, these methods usually lacked generalization, failing to produce cross-species analyses of movement and unifying frameworks for trajectory simulation. Deep learning offers a complementary paradigm: rather than encoding explicit hypotheses, the model learns movement variability implicitly from data [18], [30], constrained by what the data contains rather than by what the modeler anticipated, and critically, without losing the diversity of behaviors needed for realistic scenario evaluation.

Here, we apply the fine-tuning methodology for the first time in movement ecology by adapting DiffTraj [28], a diffusion model pre-trained on 3.5 million taxi GPS trajectories, to seabird central-place foraging trips. Our biologging dataset comprises 1,630 trajectories from four Sulidae and one Phalacrocoracidae species across six colonies in Brazil and Peru (details in Table 1). The whole dataset forms ten subsets, each corresponding to a unique colony-species pair, ranging from 14 to 447 trajectories that capture a varied range of intra- and inter-specific variation in foraging strategy. The different species also exhibit varying mobility, particularly between boobies and cormorants. Despite the domain gap, both source and target datasets are two-dimensional spatio-temporal vectors representing movement, providing a sufficient basis for transfer. The diffusion model was fine-tuned once on the complete dataset using *(colony, species)* conditioning and benchmarked against four state-of-the-art methods: a GAN for seabird CPFT simulation [19], a variational autoencoder (VAE) we specifically encoded for seabird CPFT simulation, the reference HMM for CPFT simulation [14], and a central-place iSSF, each fitted independently on each subset. The five models were evaluated on an extended framework across seven movement metrics, behavioral state inference, and spatial density analysis. We further show that conditioning the model on species and colony enables generalization to previously unseen colony-species combinations from as few as ten real-world trajectories, thereby decoupling movement simulation from data volume in a way that no current approaches achieve.

**Table 1:** Tracking datasets used in this study.

| Breeding colony | Species |  |  | Total<br>per colony |
| --- | --- | --- | --- | --- |
|  | *Sula dactylatra*<br>(Masked booby) | *Sula leucogaster*<br>(Brown booby) | *Sula sula*<br>(Red-footed booby) |  |
| Abrolhos | 98 | 38 | 0 | 136 |
| Fernando de Noronha | 187 | 14 | 48 | 249 |
| Santana | 0 | 145 | 0 | 145 |
| São Pedro e São Paulo | 0 | 447 | 0 | 447 |
| Total per species | 285 | 644 | 48 | 977 |

**Table 1:** Tracking datasets used in this study.
| Breeding colony | Species |  | Total<br>per colony |
| --- | --- | --- | --- |
|  | *Sula variegata*<br>(Peruvian booby) | *Leucocarbo bougainvillii*<br>(Guanay cormorant) |  |
| Guañape | 47 | 0 | 47 |
| Pescadores | 412 | 194 | 606 |
| Total per species | 459 | 194 | 653 |

## 2 Materials and methods

### 2.1 Data collection and preprocessing

GPS loggers (igot-U 120 and 600, Mobile Action; and AxyTrek, Technosmart) were attached to incubating or chick-rearing seabirds at colonies in Brazil and Peru (Tables 1a and 1b). Brazilian campaigns (2015–2023) covered four colonies: Abrolhos, Fernando de Noronha, Santana, and São Pedro e São Paulo Archipelagos. Peruvian campaigns (2007–2015) covered two colonies: Guanãpe and Pescadores Islands. Five species were tracked: Masked booby (*Sula dactylatra*), Brown booby (*S. leucogaster* ), Red-footed booby (*S. sula*), Peruvian booby (*S. variegata*), and Guanay cormorant (*Leucocarbo bougainvillii* ). The complete dataset contains 1,630 trajectories (977 from Brazil and 653 from Peru) organized into 10 subsets, each corresponding to a unique species-colony pair.

Each track corresponds to one complete foraging trip, with each individual contributing to one to three tracks. Original acquisition frequencies ranged from 1 second to 5 minutes, with gaps of varying frequency and duration due to advances in biologging technology and practices across campaigns. The 5-minute time step was considered sufficient to capture relevant foraging information for the five species, and thus a linear re-interpolation to a uniform 5-minute time step did not introduce excessive fake positions given the range of acquisition frequencies. Trajectories with gaps exceeding 30 minutes (6 interpolated steps) or representing more than 20% of total trip duration were excluded to preserve interpolation realism. The longest retained trajectory consists of 200 steps (16 hours 40 minutes), which matches the format used by DiffTraj.

Deep learning model training requires constant vector sizes. Consequently, all trajectories were zero-padded with stationary points at the colony location so that all time series consist of 200 steps.

### 2.2 Problem formulation

The task consists of sampling complete two-dimensional spatiotemporal trajectories **x** ∈ ℝ^*N ×*2^, where each row represents a position (longitude, latitude) at a constant time step. In the applications presented here, trajectories consist of *N* = 200 positions spaced 5 minutes apart, and **x** ∼ *q*(**x**), where *q* denotes the true data distribution. The final generative model must also be conditioned by a vector **w** = (*c, s*) representing the colony and the species.

The starting point is a model pretrained to sample from 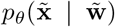, where 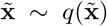 is the distribution of car trajectories and 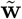 is the conditioning space of the pretrained model (e.g., total distance, average speed, origin and destination).

The goal is to fine-tune all parameters of this model on seabird trajectories, in order to sample from *p*_*θ*_(**x** | **w**), where **x** ∼ *q*(**x** | **w**) is the true conditional distribution of seabird trajectories for a given colony and species pair. Fine-tuning is performed by initializing all weights from the pretrained model and continuing training on the seabird dataset. The resulting model is evaluated by comparing generated and real trajectories across a set of metrics.

### 2.3 Diffusion model architecture and fine-tuning

#### Forward and reverse processes

Diffusion models are based on a two-way process: the *forward process* (diffusion) and the *reverse process* (denoising). The forward process iteratively adds noise to the real data over *T* computational time-steps until it becomes pure noise. The reverse process, which corresponds to the learning phase, learns to iteratively undo this noise. Through this denoising training, the model is then capable of generating new data from a noise distribution. The state-of-the-art frameworks are based on denoised diffusion probabilistic models (DDPMs) [38] [39] [40], and have outperformed Generative Adversarial Networks (GANs) and Variational Auto-Encoders (VAEs) in many domains such as computer vision [41], and matched them in others such as trajectory generation [26].

Given a trajectory sample **x**_0_ ∼ *q*(**x**_0_), the forward process is a fixed Markov chain:

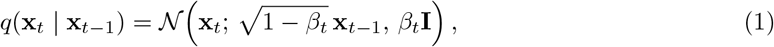

where 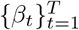 is a variance schedule and **I** is the identity matrix. Defining 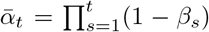, sampling **x**_*t*_ can be done at any computational time-step:

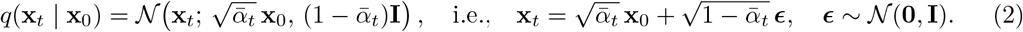

The reverse process is modeled as:

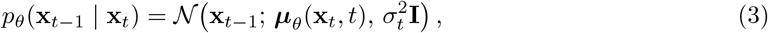

where a neural network ***ϵ***_*θ*_(**x**_*t*_, *t*) predicts the noise ***ϵ***, yielding the mean:

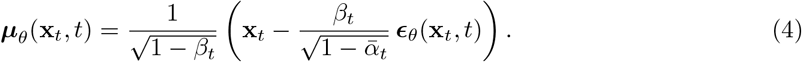

Training minimizes a simplified variational bound (a reweighted *ℓ*_2_ loss) on the predicted noise:

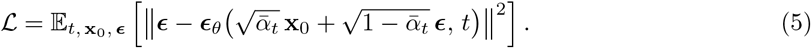

Generation is then done by iteratively sampling **x**_*t*−1_ ∼ *p*_*θ*_(**x**_*t*−1_ | **x**_*t*_) for *t* = *T, T* − 1, …, 1, starting from **x**_*T*_ ∼ *N* (**0, I**).

#### Foundation model and fine-tuning strategy

The foundation model is DiffTraj [28], pre-trained on approximately 3.5 million GPS trajectories from taxis in Chengdu, China. Each trajectory is represented as a spatio-temporal array of shape (200, 2), paired with an 8-dimensional conditioning vector. In the original model, this vector combines five continuous attributes (total distance, total duration, mean step length, mean step duration, and number of GPS points), processed by a linear layer, with three categorical attributes (departure time slot, and start and end pixel coordinates on a gridded map of the city), each processed by a dedicated embedding layer. The denoising network is a one-dimensional convolutional U-Net operating along the trajectory dimension, with four resolution levels (channel multipliers [1, 2, 2, 2], base width 128), two residual blocks per level, and self-attention at the coarsest resolution. The conditioning vector is embedded through a Wide-and-Deep module and added to the diffusion timestep embedding. The model contains approximately 16 million parameters.

For fine-tuning to seabird movement, trajectories were temporally interpolated to a constant time step such that the longest trajectory consists of 200 points, and shorter trajectories were zero-padded with stationary points at the colony after the end of the trip. The five continuous attributes and the departure-time index were set to zero, while the two remaining categorical indices were repurposed to encode the colony (values in [0, 5], six colonies) and the species (values in [0, 4], five species). This repurposing of the conditioning vector, from continuous trajectory properties to categorical ecological labels, was performed without modifying the architecture of the conditional block, demonstrating that this component can itself be re-trained toward labeled environmental conditioning, which is far more relevant for ecological simulation.

Fine-tuning was performed over 200 epochs on the combined set of 10 subsets, distinguished via conditioning, starting from the pre-trained weights. We used the Adam optimizer with a learning rate of 2 × 10^−4^, and a batch size of 32. The diffusion process used *T* = 500 steps and generation used classifier-free guidance with a guidance scale of 3. As no held-out validation set was used, given the limited size of several subsets (see Section 2.5), the epoch with the lowest training loss was retained. For comparison, DiffTraj was also trained from random initialization using the same configuration for 3000 epochs.

### 2.4 Benchmarked baseline models

#### Generative Adversarial Network (GAN)

The convolutional GAN architecture proposed by Roy et al. [19] and specifically developed for seabird CPFT simulation, was trained independently on each colony-species subset for 5000 epochs, as done in the paper. GANs are based on adversarial training between a generator and a discriminator, and are prone to instability during training and mode collapse.

#### Variational Autoencoder (VAE)

A custom VAE was implemented and trained independently per subset for 1000 epochs, as its performance did not improve beyond that point. Several architectures (fully connected, convolutional, graph convolutional) were benchmarked, and the convolutional architecture was selected based on performance. VAEs provide stable training and a continuous latent representation, generating smooth trajectories. However, they are often less able than diffusion or adversarial models to simulate real-data complexity.

#### Hidden Markov Model (HMM)

A 4-state switching HMM following Michelot et al. [14] for land-based marine predators simulation of foraging trips was fitted per subset, extending the seabird adaptation of Roy et al. [19]. The four behavioral states are interpreted as *outbound, search, forage*, and *inbound*, with the outbound and inbound states implemented as biased random walks directed away from and toward the colony, respectively. Transition probabilities depend on time-at-sea as a covariate. An incremental training of four models of increasing complexity was used to facilitate convergence. During generation, the initial orientation of each trajectory is sampled from the per-colony and per-species distribution of first steps, and the generation process stops as soon as the trajectory returns within a 2 km radius of the colony. Land points were not rejected, as this constraint does not apply universally across species and has limited impact on most evaluated metrics.

#### Integrated Step Selection Functions (iSSFs)

The iSSF framework [15] models movement as a function of habitat covariates and step kinematics. We implemented a central-place extension by adding a homing term *δ* ·*d*(*s, c*) ·*ϕ*(*t*), where *d*(*s, c*) is the distance from a candidate location to the colony, and *ϕ*(*t*) is the elapsed time, both being min-max normalized . The full model is:

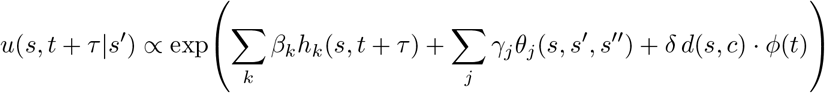

Here *u*(*s, t* + *τ* | *s*^*′*^) is the relative probability of moving to the candidate location *s* from current location *s*^*′*^ and the previous one *s*^*′′*^, after a step of fixed duration *τ* . The first sum models habitat selection, where *β*_*k*_ are the estimated coefficients and *h*_*k*_(*s, t* + *τ* ) are environmental covariates evaluated at position *s* and time *t* + *τ* . The second sum models step kinematics, where *γ*_*j*_ are estimated movement kernel coefficients and *θ*_*j*_(*s, s*^*′*^, *s*^*′′*^) are kinematic features, such as step length and turning angle. The third term is the homing penalty described above, with *δ* the learned coefficient. All environmental covariates have been standardized.

The parameters were estimated by conditional logistic regression, while the movement kernel parameters were recovered from regression coefficients using the update equations of Avgar et al. [15]. The environmental covariates included sea surface temperature, chlorophyll-*a*, eastward and northward wind, and salinity, after collinearity checks that excluded air density, wind curl and divergence, waves velocity and sea surface height. The variable selection was based on previous works on environmental covariates for seabird movement analysis identifying the oceanic fronts [42], [43], [44], [45], as well as wind importance in foraging and displacement strategies [46], [47], [48]. Simulated points on land were rejected as they could not be evaluated through habitat selection for marine variables. Of the five models, the iSSF is therefore the only one that is explicitly given the coastline and is constrained to follow it, without necessarily involving any real learning.

The environmental rasters were obtained from the Copernicus Marine Data Store (https://data.marine.copernicus.eu/products?disc=facets).

### 2.5 General training and simulation setup

Summary tables of training and trajectory generation times are available in the Supplementary A. The diffusion model was trained on the 10 subsets jointly while the number of trajectories per subset is highly unbalanced as it ranges from 14 to 447. Thus, in order to make the training balanced between subsets, the ones with fewer than 300 trajectories (i.e. all except Brown boobies in São Pedro e São Paulo and Peruvian boobies in Pescadores) were augmented by adding Gaussian noise to real trajectories until reaching 300 trajectories per subset. Noise amplitude was calibrated so that the resulting distributional error, caused by augmentation with respect to the original real data, did not exceed 5% of the lowest Earth Mover’s Distance (EMD, or Wasserstein distance) error for each metric from the subsets generated by the other models, as described below in the evaluation framework (see Methods, Section 2.6). By doing so, we included slight variability in the least numerous subset in order to make them as important as the most numerous ones.

The training and selection of deep learning models did not rely on a separation into training, test, and evaluation sets. Indeed, since the data subsets are highly heterogeneous, with some containing fewer than 50 trajectories, an evaluation based on ten or fewer trajectories would be meaningless and would deprive the training of a significant portion of the actual distribution. The absence of overfitting is therefore not guaranteed for the GAN and the VAE, and is evaluated for the diffusion model via conditioning through leave-one-out (Section 3.3). HMM and iSSF models were trained on original (non-padded) interpolated tracks.

After training, each model generated 10 times the number of trajectories observed in the real datasets. For the diffusion model, generation occurred jointly across all species and sites using label-based conditioning. The other models produced data subset by subset. For deep learning models, trained on padded data, the generated trajectories were un-padded by cutting them when more than 4 positions in a row are simulated within a common radius of 0.5 km, while being less than 2 times as far from the colony as the 200^*th*^ generated position. This allows robust un-padding, since the trajectory scales vary from a subset to the other and no single cutoff distance would suit for all subsets.

### 2.6 Evaluation framework

All evaluations were performed on the 10 independently simulated sets of 10 × *n*_real_ trajectories per model, and results are reported as mean ± s.d. to account for simulation variability.

#### Trajectory-level metrics

Seven metrics were computed for each trajectory un-padded, with five being at the whole trajectory scale and the last two incorporating the local step-scale:

1. **Maximum distance from colony**, *D*_max_
2. **Total distance traveled**, *D*_tot_
3. **Straightness index**,

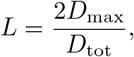

which ranges from 0 for highly non-linear trajectories to 1 for perfectly linear out-and-back foraging trips (can be greater than 1 for non-looping trajectories)
4. **Global orientation** (angle between geographic north and the line connecting the colony to the farthest point of the trajectory)
5. **Total duration**,
6. **Step length** (distance between two consecutive points)
7. **Turning angle** (angular deviation between successive steps, calculated from three consecutive points).

The trajectory-scale metrics give one value per trajectory, while the step-scale metrics are calculated on each step of the trajectory. However, the real and simulated distributions are obtained by combining the values of all trajectories for both metric scales.

#### Distributional comparison and global scores

For each metric *m*, the EMD between the generated distribution 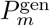 and real distribution 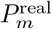 of all trajectories was computed:

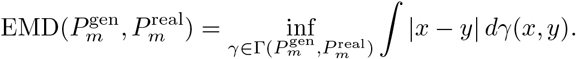

For the global orientation, the bi-dimensional Wasserstein distance was applied.

To normalize and compare the error made across the seven different metrics, distributions were standardized using the mean and standard deviation of the real data for each metric. The global error score which combines the seven metrics is:

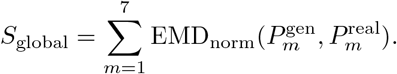

A global diversity error was also computed as

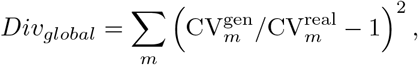

where CV = *std/mean* is the coefficient of variation, assessing whether generated diversity matches real diversity (penalizing both mode collapse and over-dispersion).

#### Behavioral state inference

A 3-state HMM (travel, search, forage) was fitted per subset on real trajectories. The Viterbi algorithm [49] was then used to infer the most likely state sequence for each real and simulated trajectory. State proportions, temporal state distributions over trajectory progress, and empirical state transition matrices were compared between real and generated data for each model.

#### Spatial distribution analysis

Foraging positions inferred from the behavioral state inference for real and generated trajectories, and located more than 10 km from the colony were extracted. Spatial foraging intensity was estimated using kernel density estimation (KDE).

#### Conditioning sensibility

A leave-one-out protocol was used to evaluate generalization of the diffusion model. The diffusion model, already pre-trained on vehicle trajectories, was further fine-tuned on the nine subsets excluding Brown boobies at Santana (*n* = 145). This subset represents a known species in an unknown colony. The resulting zero-shot model was then evaluated on the held-out subset, and progressively fine-tuned during 200 epochs using 10, 20, 30, 50, 80, 110, and 145 trajectories from Santana. Trajectory selection within the subset was random, but with a fixed seed. Thus, the 20 trajectories include the first 10, and so on. The learning rate was divided by 10 with respect to the one used on the nine subsets to avoid overfitting on the new subset. Each model obtained from fine-tuning run sampled 1450 trajectories. Each generated set of 145 trajectories was evaluated against the full real set. A reference baseline was established by evaluating the random real subsamples used at each size.

## 3 Results

We evaluated five classes of generative models on ten colony-species trajectory subsets. Results are organized into three sections: (i) an overall benchmark of the five models, (ii) the dynamics of fine-tuning convergence, and (iii) the contributions of conditioning to generalizability and data efficiency. Since the results cover three aspects of model evaluation, each part is illustrated on a representative subset, selected to best reflect the process under evaluation. The full results across all colony-species subsets are provided in Supplementary Information B. To account for simulation variability, each model generated ten times the number of real trajectories. The trajectory visualizations only show a single simulated set, matching the size of the real dataset, while the statistical analyses are performed on the inflated simulated datasets.

### 3.1 Diffusion model outperforms all baselines across all movement scales

The comparison of models’ performance is illustrated here using the Peruvian booby (*Sula variegata*) at Pescadores, Peru (*n* = 412, representing 25% of the full dataset), which is the second subset with the most trajectories and is therefore, in principle, best described by real data.

#### Visual and qualitative assessment

A visual comparison of the 412 trajectories of Peruvian boobies at the Peruvian site of Pescadores Island generated by each model (Fig. 2) reveals the typical characteristics of the model classes. The GAN produces trajectories that are individually realistic but, collectively, underrepresent the actual spatial diversity. Long, northward-bound trajectories are not reproduced. The VAE simulates smoothed trajectories that remain close to the colony, reproducing an average movement with significantly reduced spatial range and diversity. The HMM generates spatially highly dispersed trajectories without directional persistence and does not reproduce any large-scale spatial realism related to movement at sea. The iSSF generates trajectories highly concentrated around the colony due to the strong central-place homing term, and produces unrealistic explorations at sea, particularly southward. The diffusion model simulates trajectories that most closely resemble the actual distribution in terms of spatial exploration, coastal orientation, and diversity of shapes. This realism holds for all ten subsets (see Supplementary Information B for other subsets’ visualization). The Guanay cormorant (*Leucocarbo bougainvillii* ) at Pescadores, the only non-Sulidae species in the dataset, exhibits markedly shorter trips and a more coastally constrained spatial range than the boobies, reflecting its wing-propelled diving strategy and higher energetic cost of flight (Supplementary Fig. 19). The diffusion model reproduces this species-specific movement envelope with the same fidelity observed across booby subsets.

**Fig. 1:**
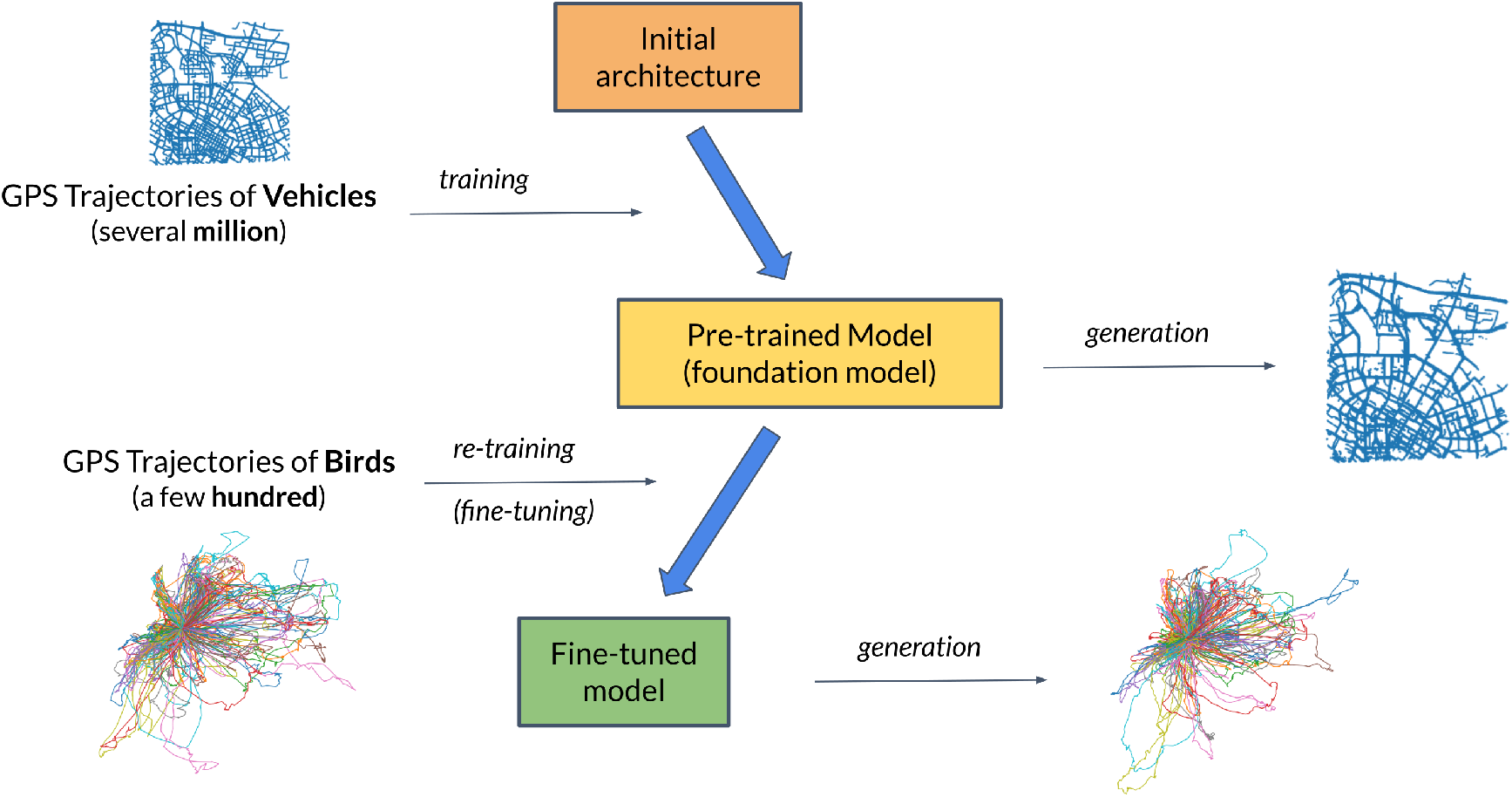
Transfer learning framework for seabird trajectory generation. From top to bottom: the initial architecture, here a diffusion model [28], is first pre-trained on several million GPS trajectories of vehicles, learning general spatiotemporal movement patterns. The resulting foundation model can simulate realistic vehicle trajectories. By fine-tuning it on a few hundred seabird GPS tracking trips, the model adapts its learned representations of trajectories to the ecological context of central-place foraging trips. The fine-tuned model becomes able to simulate ecologically realistic seabird trajectories.

**Fig. 2:**
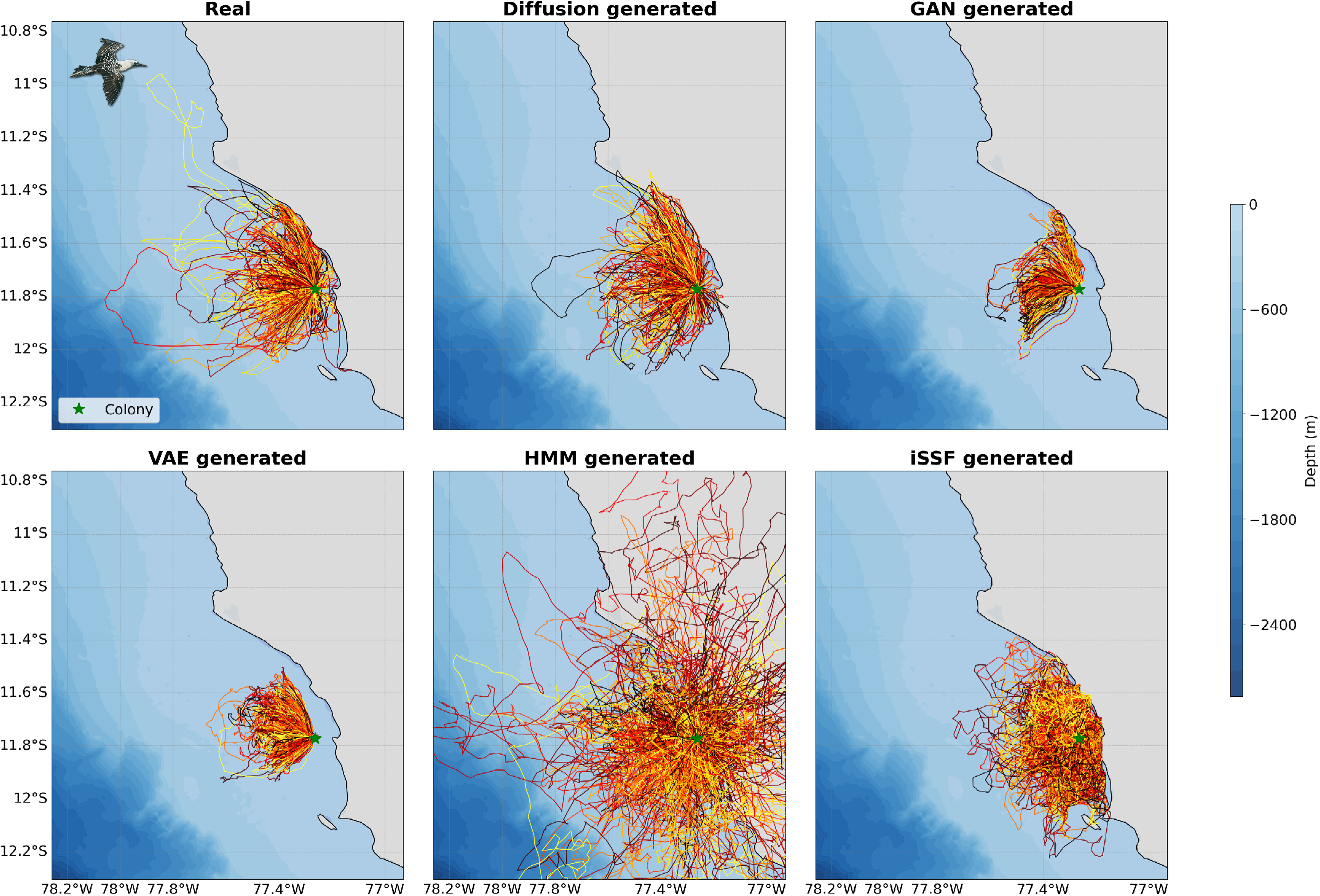
Real and model-generated trajectories for Peruvian boobies (*Sula variegata* ) at Pescadores Island, Peru. Each panel shows 412 trajectories. A trajectory is an out-and-back trip from the colony. Colors represent unique trips and are randomized for visual clarity.

#### Quantitative metric analysis

The distribution of the seven movement metrics (Figs. 3 and 4) quantitatively confirms the superior performance of the diffusion model. Notably:

- Step length distribution: the actual distribution is bimodal, reflecting different behaviors associated with travel (long steps) and foraging (short steps). Diffusion and the HMM are the only models to reproduce this bimodal or even trimodal distribution (when considering a search state with intermediate steps), although the HMM is explicitly trained on the distributions of step length and turning angle. The GAN and VAE struggle to reproduce this bimodality, while the iSSF reproduces only a single average distribution.
- Straightness index: this metric *L* = 2*D*_max_*/D*_tot_ captures the joint relationship between the maximum distance from colony, *D*_*max*_, and the total distance of the trajectory, *D*_*tot*_. It ranges from 0 (highly non-linear trajectories) to 1 (perfectly linear out-and-back, see Methods 2), and captures a trajectory-scale geometry. The diffusion model most realistically reproduces the actual distribution, concentrated around 0.8-0.95, reflecting the relatively straight flight paths of Peruvian boobies, while the GAN and VAE under- and over-estimate it, respectively. The HMM and iSSF show no realism, although HMM distributions of *D*_*max*_ and *D*_*tot*_ are not that far from the real ones, while the iSSF dramatically over-estimates *D*_*tot*_.
- Overall, the diffusion model realistically reproduces all real distributions, with a slight smoothing effect. In contrast, the GAN and the VAE struggle to achieve realism without over-representing certain parts of the distributions, resulting in an imbalance between the generated distributions and the real ones. Finally, the HMM and the iSSF, trained at the step scale, faithfully reproduce local metrics, with the caveat that the iSSF considers only unimodal distributions.

**Fig. 3:**
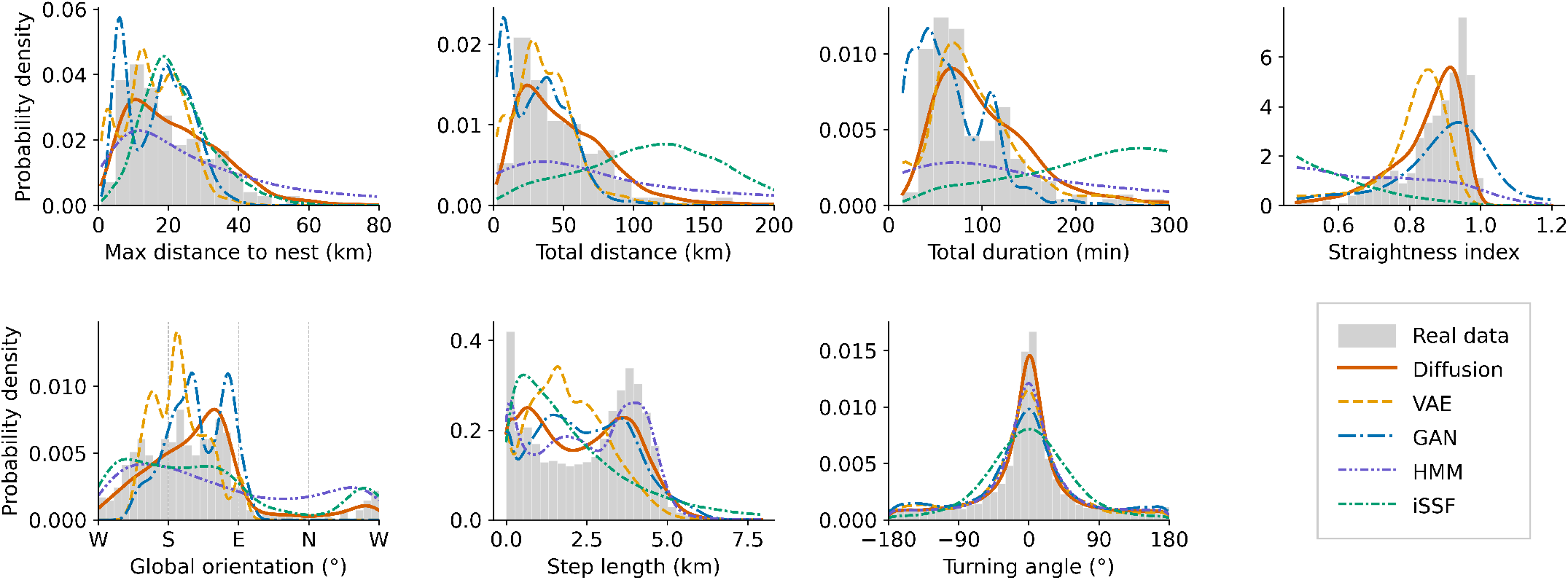
Distributions of seven movement metrics for real and model-generated trajectories of Peruvian Boobies (*Sula variegata* ) at Pescadores Island, Peru (*n* = 412). Distributions are shown for real trajectories (gray) and those generated by five models: diffusion (red), VAE (yellow), GAN (blue), HMM (purple), and iSSF (green). For each model and each metric, the curves represent the probability density of the generated values, estimated using Kernel Density Estimation (KDE) with a Gaussian kernel. As each model generated ten times the number of real trajectories, the 4,120 values produced by a given model are treated as a continuous distribution and smoothed into a single KDE curve for direct comparison with the histogram of the real data.

**Fig. 4:**
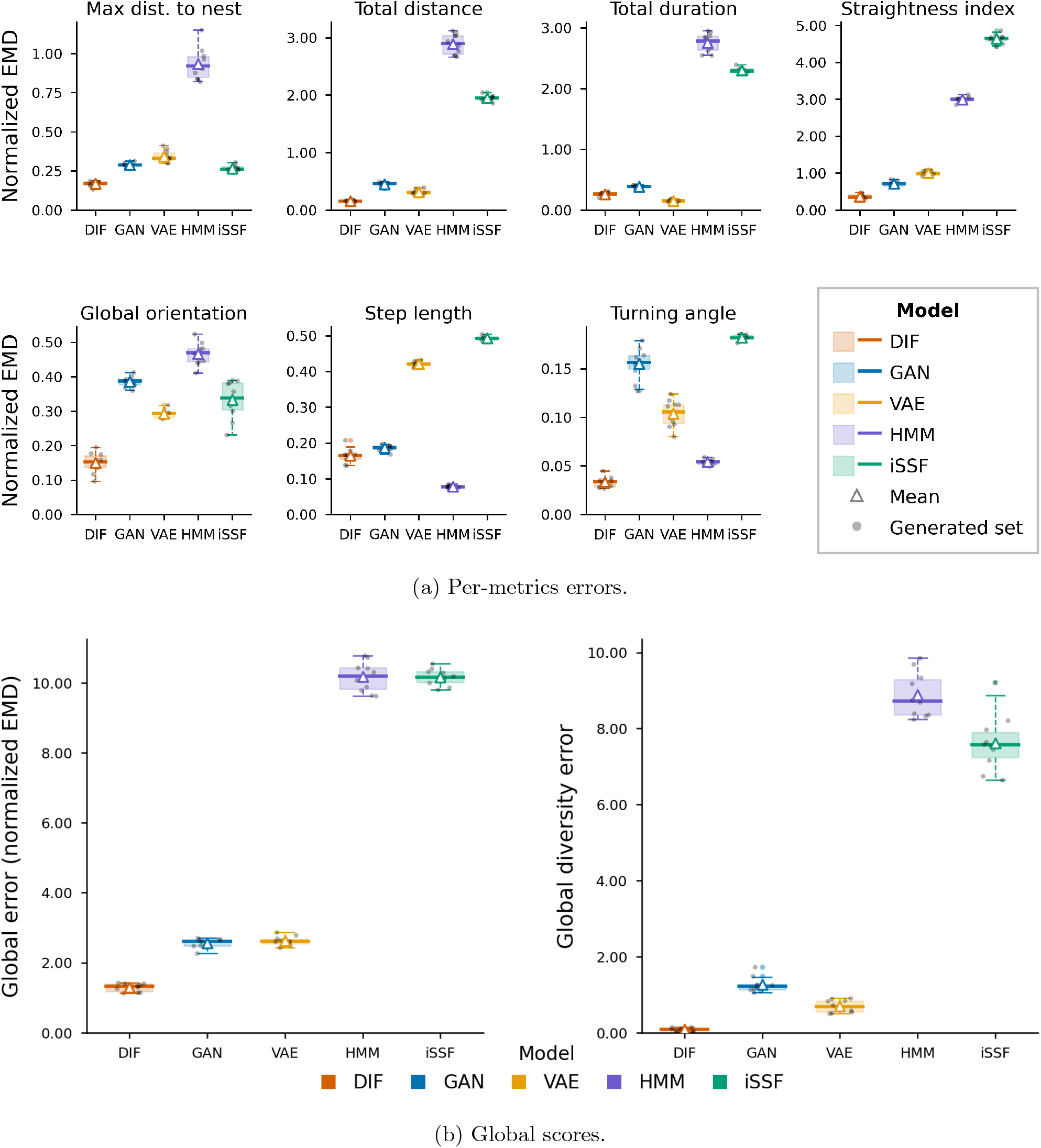
Per-metric and global error scores comparing real and model-generated trajectories for Peruvian Boobies (*Sula variegata* ) at Pescadores Island, Peru (*n* = 412). Panels show a) per-metrics errors, b) global errors and global diversity errors. Errors were obtained by applying the Earth Mover’s Distance (EMD, or Wasserstein distance) between real and generated distributions. For each model, boxplots were obtained from 10 independently generated sets, each evaluated against the full real dataset.

The global error scores and metrics via boxplots (Fig. 4) confirm that the diffusion model achieves the lowest errors across all subsets of the data (see Supplementary Information B for all results), including in terms of the diversity generated. Unlike other models, which exhibit significant weaknesses for some metrics, often distinguishable by the scale considered, the diffusion model captures both local and global dynamics and outperforms the best of the models at every scale.

#### Behavioral state analysis and spatial distribution

To assess the behavioral realism of generated trajectories, we fitted a 3-state HMM to the real trajectories of Peruvian boobies from Pescadores, Peru (emission distributions in Supplementary Fig. 22)), and used the Viterbi algorithm to infer behavioral states on both real and generated trajectories from all five models. The three states are interpreted as *traveling, searching*, and *foraging* based on their step length and turning angle distributions (Fig. 5a). Each model generated ten times the number of real trajectories (4,120 trajectories in total), and state inference was applied identically across all datasets.

**Fig. 5:**
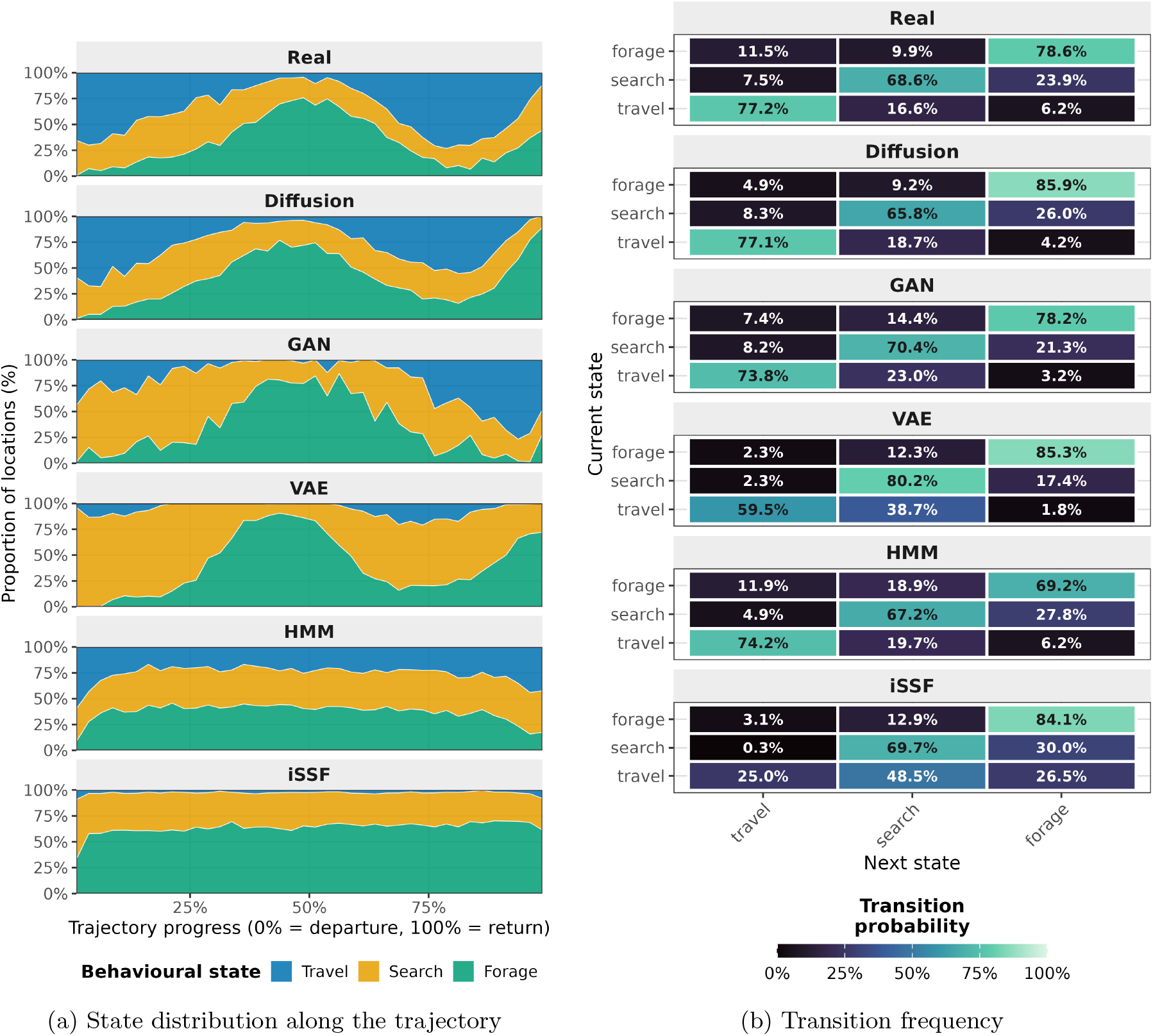
Behavioral state dynamics for Peruvian boobies (*Sula variegata* ) from Pescadores Island, Peru. States were assigned using a 3-state Hidden Markov Model (travel, search, forage) fitted on real trajectories and applied to all datasets (emission distributions and state proportions in Supplementary Fig. 22). **a)** Distribution of behavioral states along normalized trajectory progress (0% = departure, 100% = return) for real trajectories and the five generative models (Diffusion, GAN, VAE, HMM, iSSF). **b)** Transition probability matrices between behavioral states in each model.

Behavioral analysis via HMM inference (Fig. 5) shows that the diffusion model reproduces the behavioral sequence of a Peruvian booby CPFT at Pescadores: primarily travel during the initial and final phases of the trajectory, while foraging and searching occupy the middle. Upon approaching the colony on the return trip, the shorter step-length segments identified as search and foraging states, reflecting peri-colony approach behavior and rafting, are accurately reproduced in both proportion and temporal placement. The proportions of states in the trajectories generated by diffusion (traveling 30%, searching 30%, foraging 40%) are close to the actual proportions (traveling 38%, searching 30%, foraging 33%), with foraging slightly overrepresented at the expense of travel. The GAN reproduces the proportions reasonably well but with a more erratic temporal structure. The VAE produces almost no travel and overrepresents the intermediate search state. The HMM does not reproduce behavioral temporality, specifically the sequence of search followed by foraging, despite realistic state proportions and transition probabilities. The iSSF dramatically overrepresents foraging and avoids travel, consistent with its spatial over-concentration observed in Fig. 2.

The spatial distribution of inferred foraging points (Supplementary Information C), estimated using kernel density estimation (KDE), reveals that the diffusion-generated distribution replicates the spatial asymmetry concentrated toward the north and west of the foraging intensity characteristic of Peruvian boobies at Pescadores, notably with the secondary hotspot on the coast north of the colony. The GAN and VAE produce foraging states concentrated around the colony without spatial exploration. The HMM produces a more dispersed distribution but with no realistic direction other than that imposed by the starting angle provided to the simulation. The iSSF is spatially over-constrained around the colony, with no offshore orientation.

### 3.2 Cross-domain transfer enables rapid adaptation to seabird movement

We evaluate the model’s adaptation speed during fine-tuning when transitioning from car to seabird trajectories, illustrated on Masked boobies (*Sula dactylatra*) at Abrolhos, Brazil (*n* = 98), a subset chosen because it combines a wide directional diversity, a broad range of trip distances and durations, and a strong return-to-colony structure: characteristics that maximally contrast with car trajectories and thus constitute a demanding test of domain adaptation (Fig. 6). At epoch 0, the foundation model generates trajectories characteristic of urban vehicle movement: spatially compact, following straight grid-like paths without any circular return structure. After a single pass of the seabird dataset (epoch 1), the model has already adjusted to a biologically realistic spatial scale of hundreds of kilometers, producing curvilinear paths that no longer follow road-like routes and beginning to reproduce a return-to-colony structure. By epoch 2, the generated trajectories begin to be indistinguishable from real ones in terms of scale, shape diversity, and colony return, an adaptation achieved in approximately 30 seconds of training. Epochs 3 to 200 yield gradual multi-scale refinement while maintaining realistic diversity. This rapid convergence reflects the transferability of two priors learned from car trajectories: a representation of smooth two-dimensional spatial sequences, and an implicit understanding of departure-displacement-return structure that generalizes across domains.

**Fig. 6:**
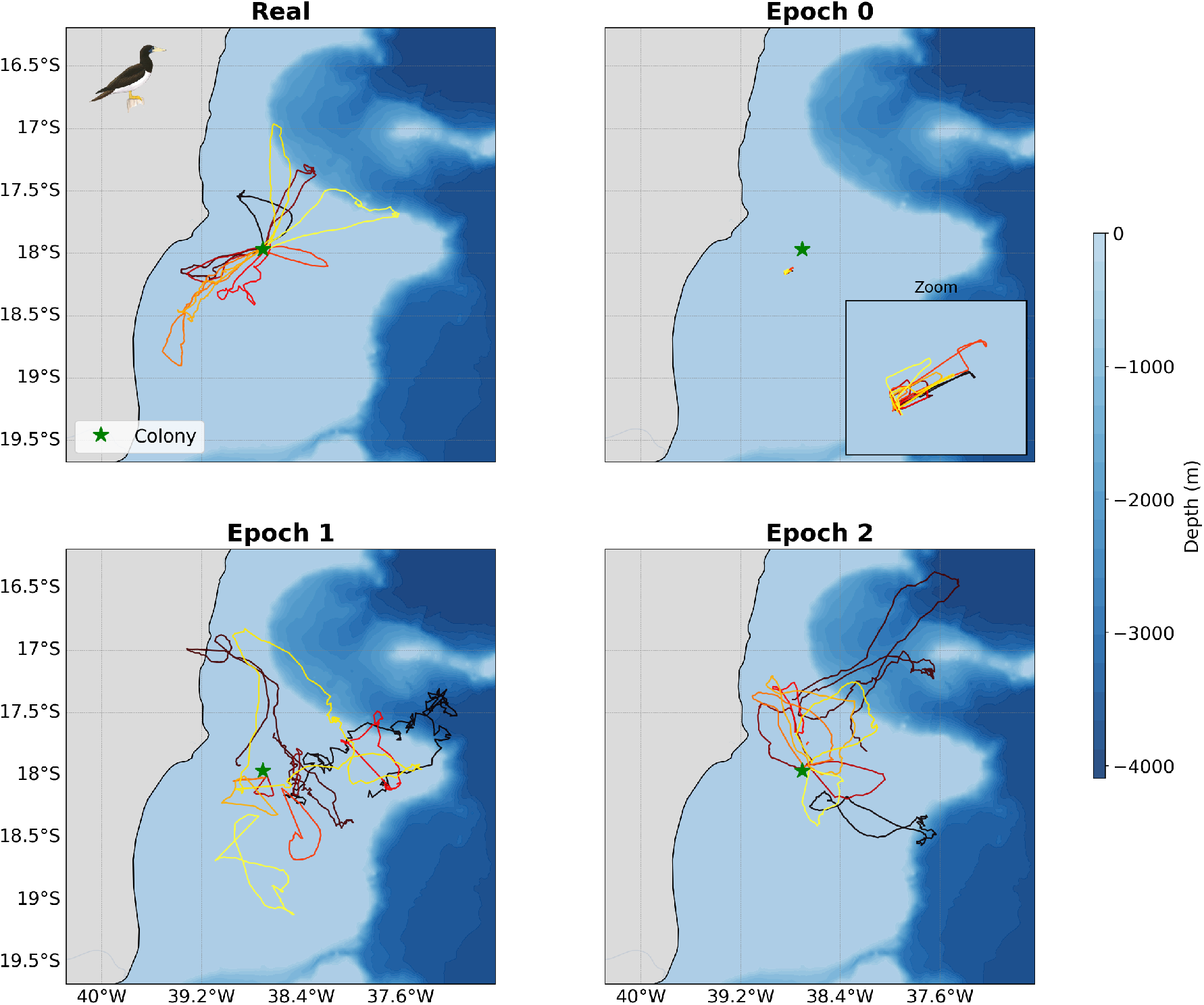
Real and model-generated trajectories of Masked Boobies (*Sula dactylatra* ) at Abrolhos Archipelago, Brazil, across the first epochs of fine-tuning. Each panel shows 10 trajectories. Generated trajectories are sampled at successive epochs of fine-tuning (where one epoch corresponds to a single full pass of the training dataset through the model), with epoch 0 corresponding to the foundation model pre-trained on car trajectories. Fine-tuning was implemented on the complete dataset jointly through conditioning and is illustrated here for this species-colony subset.

To achieve equivalent quantitative performance from random initialization, DiffTraj required approximately 3000 epochs (7 hours, 14 times longer; all training and simulation durations are reported in Supplementary Tables 1 and 2).

### 3.3 Intra-domain conditioning enables zero-shot generalization and few-shot adaptation

We trained the diffusion model on nine subsets, and then iteratively re-trained it on the tenth subset alone. We chose to leave out the subset of Brown boobies (*Sula leucogaster* ) at Santana Archipelago, Brazil, since this species occurs in other colonies, whereas Santana hosts no other species in the dataset, making it a rigorous test of spatial generalization. The model fine-tuned on the nine subsets is already capable of generating realistic trajectories for Brown boobies at Santana with zero target-domain data (Fig. 7, values at 0 trajectories), with global and metric errors already among the best across all models evaluated on the full dataset (Supplementary Fig. 12). This shows that having seen Brown boobies in other colonies is sufficient to simulate their behavior with high realism in an unknown location. As trajectories from Santana are progressively added, performance improves immediately (Fig. 7). After seeing just 10 of the 145 available trajectories, the model already outperforms all baselines trained on the full dataset (Supplementary Fig. 12). The global orientation, which is the most colony-specific metric as it reflects local geography and oceanography, is adjusted from the very first trajectories. After 30 trajectories, the model reaches near-optimum performance across all metrics. Furthermore, between 10 and 50 trajectories, the model better represents the full distribution of 145 real trajectories for certain metrics than the real subsample itself provided for training, generating trajectories that exceed the coverage of the training subset alone.

**Fig. 7:**
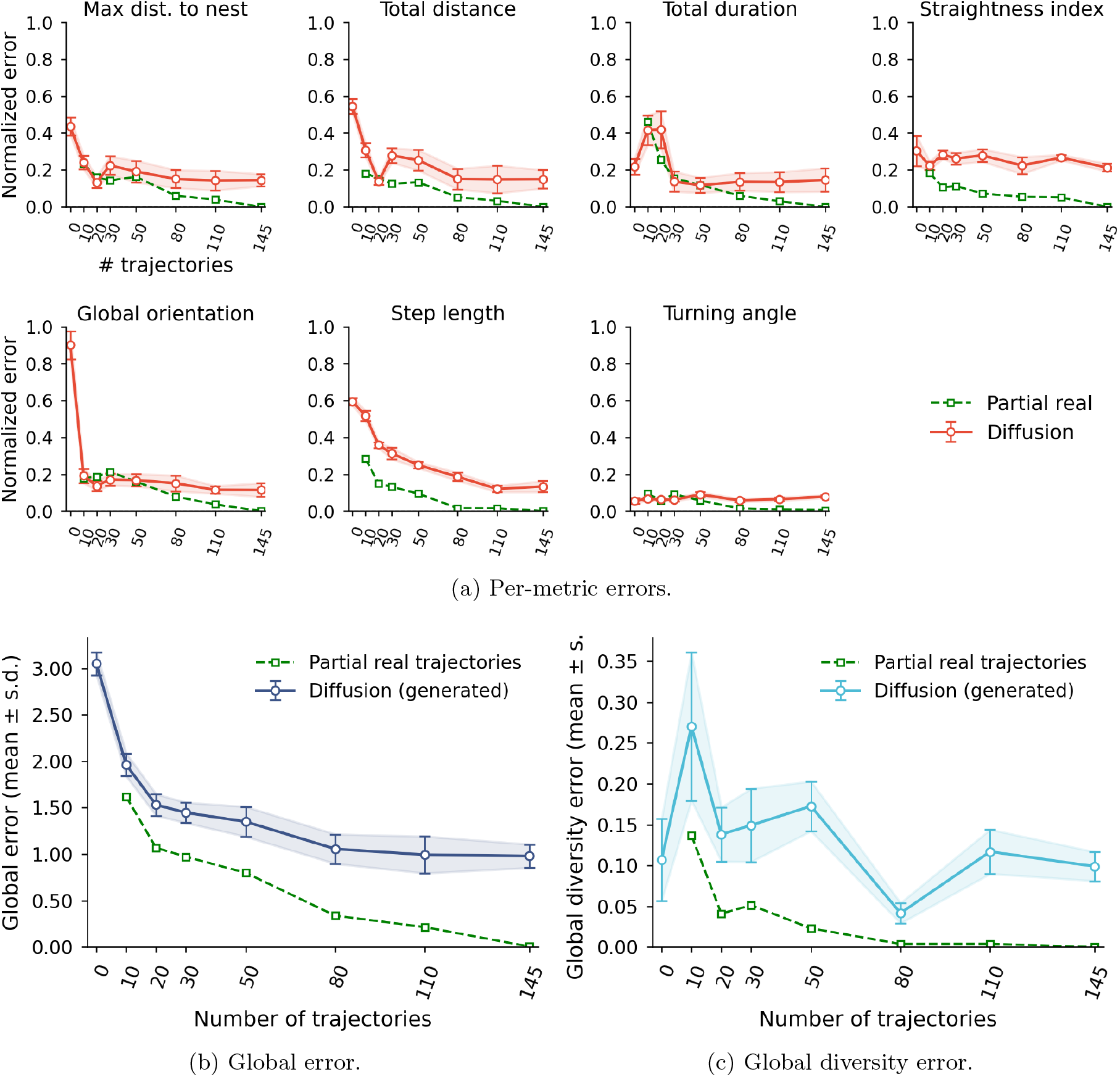
Few-shot adaptation performance of the diffusion model for Brown boobies (*Sula leucogaster* ) at Santana Archipelago, Brazil, as a function of the number of trajectories used for fine-tuning. For each subset size, DiffTraj was fine-tuned from a foundation model pre-trained on the 9 remaining subsets, then used to generate 145 synthetic trajectories across 10 independent runs, each evaluated against the full set of 145 real trajectories (mean ± s.d.). The dashed curves show scores obtained from an equivalent random subsample of real trajectories, representing the baseline error attributable to data scarcity alone. Metrics in **a)** are normalized so that a score of 0 indicates perfect agreement with the real distribution. Per-metric heatmaps are provided in Supplementary Fig. 24.

## 4 Discussion

The capacity of a model pre-trained on urban vehicle trajectories [28] to outperform purpose-built seabird movement models across all evaluated metrics demonstrates that trajectory-scale structure is a transferable property that transcends ecological and geometric domain boundaries.

Treating the complete trajectory rather than a sequence of steps allows us to account for long-term dependencies that step-by-step movement models cannot represent. This limitation is structural: the Markovian construction of both HMMs and iSSFs prevents them from reproducing large-scale trajectory complexity even when a return constraint is imposed, a known limitation of these methods [14], [19], [9]. The HMM’s failure to reproduce behavioral temporality despite realistic state proportions and transition probabilities illustrates this structural ceiling directly: the Markovian assumption prevents the model from capturing the sequential organization of foraging and searching across the full trip duration. The iSSF’s dramatic overrepresentation of foraging states and spatial over-concentration around the colony are likewise structural: the framework fits a single step-length and turning angle distribution across all behavioral contexts, and the homing penalty continuously attracts simulated locations toward the colony as elapsed time increases, suppressing offshore exploration even when habitat covariates would otherwise support it. The straightness index is particularly revealing, as birds that travel farther tend to fly more directly, reflecting an energy-efficiency trade-off between distance covered and flight path tortuosity. This correlation between maximum distance from the nest and total trip length is correctly reproduced only by deep learning models, and with high fidelity only by the diffusion model. This overall consistency does not come at the expense of local precision: the distributions of step length and turning angle and their multimodality are also reproduced with greater precision by the diffusion model than by the other models, despite the explicit step-by-step formulation of HMMs and iSSFs. The approach used here by deep learning models, by considering the trajectory in its entirety, can implicitly learn the set of global constraints without having to encode them explicitly [26], [27]. This capacity to represent the trajectory globally is precisely what realistic foraging behavior demands: a central-place foraging trip integrates decisions across multiple spatial and temporal scales, from fine-grained search movements to the overall outbound-and-return geometry, a multi-scale structure that step-by-step models cannot hold together.

Several factors account for this performance gap between the three deep learning models. First, diffusion models are known for being more stable during training and more robust to overfitting than GANs and VAEs, while producing varied outputs without the mode collapse or hallucinations that often plague these architectures [50], [51]. In our case, the GAN fell into mode collapse, reproducing a restricted subset of trajectory shapes with little variation and failing to capture the northward long-distance foraging trips characteristic of Pescadores Peruvian boobies. Second, DiffTraj consists of approximately 16 million parameters, whereas the GAN has about 1 million and the VAE has 200,000, giving it a much greater capacity to capture the multi-scale complexity and variety of foraging behaviors. Third, DiffTraj is used here fine-tuned from a pre-training rather than a random initialization of its parameters, giving it prior knowledge of the trajectory space that the other models must construct from scratch. However, each of the approaches evaluated here remains entirely relevant depending on the context. GANs train a discriminator that can serve as an independent evaluation metric. The VAEs provide a compact latent representation for clustering and dimension reduction [50], [51]. The HMMs are very useful for behavioral state inference [52], [53], as we used in the evaluation framework. iSSFs provide valuable correlations between movement and environmental gradients [54], [16]. The superiority of this diffusion model for the realism of simulations does not replace these tools, but complements and extends this methodological palette.

Fine-tuning-based learning, which generates realistic trajectories after as few as two training epochs, along with generalization to a new colony after as few as 10 real trajectories, opens up significant opportunities for frugal modeling while advancing the state-of-the-art. Biologging campaigns are costly in terms of both money and time, while being subject to ethical constraints, making large datasets rare, particularly for remote locations and hard-to-access species [3], [4]. Being able to start training with trajectories other than those of animals, such as the widely available trajectories of millions of vehicles, and then better represent animal trajectories can change the approach to biologging campaigns, such as adjusting the sampling effort. This result aligns with transfer learning theory: both vehicle trajectories and those of seabirds are two-dimensional vectors representing a spatiotemporal sequence governed by an energy constraint, a starting point, and an endpoint [36], [37]. This common foundation is sufficient for knowledge transfer despite the wide domain gap. Car trajectories are indeed much more geometrically constrained: they follow road networks consisting of straight lines and right angles, as well as stationary points, whereas seabirds move without such constraints over the ocean. Transferability therefore occurs by relaxing constraints rather than replacing them. Although ship trajectories might appear geometrically closer to those of seabirds, models trained on them are designed for next-step prediction in collision avoidance or traffic optimization [27], [55] rather than whole-trajectory generation. DiffTraj, by contrast, was developed to generate complete trajectories for anonymization purposes [28], making its architecture directly transferable to our generative objective and a more appropriate foundation than a geometrically closer but functionally mismatched alternative.

Training all species and colonies simultaneously through conditioning is a key architectural choice. Conditioning acts as a generalization mechanism, while training independent models for each subset would mean forgoing a common representation of trajectories, species, and colonies, and would require a larger number of trajectories for each of these. Through conditioning we enabled a learning that is both specific to each subset and generalizable, transferable to new colonies. Without having observed a species’ behavior in a new location, the model, having seen it in other places, already knows how to represent realistic behaviors, though individual-level drivers of site fidelity such as memory and social information [56] remain implicitly absorbed into the population-level distribution rather than explicitly modeled. Model learning is not mere memorization of the training subsets, but an accurate and transferable representation of movement that makes extrapolation to new contexts possible.

The current framework, while effective, is a direct reuse of the architecture developed for vehicles without any modifications. A future improvement would be to implement a conditional block designed and developed specifically for ecological data. The evaluation framework considers individual trajectories and evaluates the population as the sum of individuals. However, representing a population requires taking into account its composition, the aggregate of individual variations, and interactions [57], [58], [59]. This limitation stems in part from the available data, which represents only a sample of the total population, as tagging all individuals would often be impossible, too costly, and ethically problematic [60].

The next most direct and relevant step is environmental conditioning, which enables the linking of trajectory simulations to oceanographic and habitat variables that are explicit in both space and time. This would facilitate scenario analysis, such as assessing how species respond to variations in temperature and chlorophyll-*a* distribution for each colony [61], [42], [62]. Such a conditioned model would also improve the evaluation of the impact of human-induced projects on the movement of these seabirds, such as Marine Protected Areas [63] or offshore wind farms [64], [65], allowing assessment the viability of entire colonies [66], and informing fisheries management through the use of seabirds as ecosystem indicators [24]. Conditioning by individual traits (sex, breeding stage, body condition, personality) would enable the assessment of demographic heterogeneity in foraging behavior [67], [58], and would be straightforward through labeling as done here for colony and species.

A practical consideration for ecologists wishing to apply this method concerns the minimum required dataset size. The model imposes no minimum constraint on the number of trajectories and can be trained on very few: here, subsets range from 14 to 447 trajectories, and the model was also tested on as few as 5 trajectories, producing outputs similar to those given as input. The fundamental constraint therefore does not stem from the model’s capacity, but from the representativeness of the available data: a model trained on a small or biased dataset will faithfully reproduce it, but not necessarily the population [2], [7]. The results of rapid learning via conditioning show that the model can achieve simulations surpassing other methods in terms of realism on the full dataset without having seen it entirely. However, the simulated trajectories should only be interpreted in light of the diversity and representativeness of the training dataset. One technical constraint is inherited from the foundation model: trajectories are represented as fixed-length sequences (here 200 positions), jointly constraining temporal resolution and maximum trip duration. However, because the model’s architecture is fully convolutional along the trajectory dimension (see Methods, Section 2.3), this length can be adjusted when fine-tuning to accommodate longer or finer-resolution trajectories, making it a preprocessing choice rather than a fundamental limitation.

It follows that the sampling effort required to train a reliable model scales with the ecological variability of the population: colonies or species with highly variable foraging strategies, driven by individual specialization, environmental heterogeneity, or seasonal variation, will require more extensive datasets than those with homogeneous movement patterns. The method should therefore be seen as a tool that amplifies the information contained in available data, not as a substitute for adequate field sampling.

This methodology is not specific to seabirds or CPFTs. Any species for which movement data exists that allows trajectories to be constructed can be studied and simulated using this approach. These results establish a new paradigm in movement ecology: harnessing the vast resources of trajectory data from other fields to overcome the fundamental constraint of data scarcity in animal movement research, whilst remaining computationally frugal thanks to open-source models [1], [2], [32]. As biologging datasets expand and transfer learning continues to mature, this new intersection between movement ecology and generative AI offers a powerful new platform for scenario simulation under environmental change, data augmentation for undersampled species, and, crucially, conservation planning for already threatened populations.

### 4.1 Implementation details

Deep learning models were implemented in Python using *Pytorch*. The Python version used is 3.12.2. HMM for generation were fitted using the *momentuHMM* R package [52], while the ones for state inference were implemented with the *moveHMM* R package [68]. The iSSFs were trained using the *amt* R package [16]. The R version used is 4.5.3. All models were trained on a local computer equipped with 32 GB of RAM and GPU (8 GB VRAM), the GPU being activated only for deep learning models.

## Acknowledgements

The authors wish to thank the following participants in the fieldwork campaigns that provided the datasets used in this study: Yann Tremblay, Henri Weimerskirch, Patrícia Mancini, Tatiane Pereira Xavier Nascimento, Mariano Valverde, Jaime Silva, Ainhoa Lezama, Delia Vega, Sofia Rivadeneyra, Jose Carlos Torres Maita, Claudia Arbulu, Daniel Grados, Marie Wach, Claire Saraux, Stephane Bertani, Vilma Romero, Jeremy Masbou, Wendy Flores, Carlos Calvo, Maite Aranguenã-Proaño, Leila Figueiredo, Samantha Cox, Luísa Bertolini, Cynthia Campolina, Victor Libardi and Maicon Pegoraro. We thank Clara Lerebourg for her contribution to the visual design of the figures and valuable insights on the manuscript.

## Declarations

- **Funding:** Julien Patras was funded by an AMX fellowship from École Polytechnique. This study was financed in part by the Coordenação de Aperfeiçoamento de Pessoal de Nível Superior – Brasil (CAPES) – Finance Code 001.
- **Competing interests:** The authors declare no competing interests.
- **Ethics approval:** Data collection was conducted with the approval of the Brazilian Ministry of the Environment-Instituto Chico Mendes de Conservação da Biodiversidade (permits nos. 52583 and 64261) and guidelines of Ethical Animal Use (CEUA-FURG 23116.001336/2020-40, CEUA - UFRGS 37905), and the Peruvian federal agency, the Programa de Desarrollo Productivo Agrario Rural (AGRORURAL).
- **Data availability:** Tracking datasets are available on the Seabird Tracking Database (http://www.seabirdtracking.org). Datasets not yet deposited are being prepared for submission by the respective data providers and can be requested from the corresponding author in the meantime.
- **Code availability:** The code is available on Zenodo (https://doi.org/10.5281/zenodo.21364538). As the full dataset cannot be shared, the public repository will include a toy (synthetic) dataset for demonstration purposes. The diffusion model builds on DiffTraj [28] and the GAN and HMM baselines on [19]. Raw data preprocessing was performed using the *cpforager* Python package (https://github.com/AdrienBrunel/cpforager).
- **Author contributions:** J.P. conceived and led the project, developed the code, and wrote the manuscript. R.F. contributed to project design, provided supervision, and contributed to the manuscript. A.B. performed data processing and organization and contributed to the manuscript. A.R. contributed to ideas, data collection, and the manuscript. S.B. contributed to ideas and the manuscript. S.L. conceived the project, contributed to supervision, ideas, data collection, and the manuscript. L.B., G.T.N., C.B., J.J., and G.P. contributed to data collection and reviewed the manuscript. K.D. contributed to data collection.

## References

[1] Gomez, S. et al. Understanding and predicting animal movements and distributions in the Anthropocene. Journal of Animal Ecology 1365–2656.70040 (2025). URL https://besjournals.onlinelibrary.wiley.com/doi/10.1111/1365-2656.70040.

[2] Nathan, R. et al. Big-data approaches lead to an increased understanding of the ecology of animal movement. Science 375, eabg1780 (2022). URL https://www.science.org/doi/10.1126/science.abg1780.

[3] Wilmers, C. C. et al. The golden age of bio-logging: how animal-borne sensors are advancing the frontiers of ecology. Ecology 96, 1741–1753 (2015). URL https://onlinelibrary.wiley.com/doi/abs/10.1890/14-1401.1. xeprint: https://onlinelibrary.wiley.com/doi/pdf/10.1890/14-1401.1.

[4] Watanabe, Y. Y. & Papastamatiou, Y. P. Biologging and biotelemetry: tools for understanding the lives and environments of marine animals. Annual Review of Animal Biosciences 11, 247–267 (2023). URL https://www.annualreviews.org/content/journals/10.1146/annurev-animal-050322-073657.

[5] Tremblay, Y. & Bertrand, S. in “Bio-logging” as a Tool to Study and Monitor Marine Ecosystems, or How to Spy on Sea Creatures 1 edn, (eds Monaco, A. & Prouzet, P.) Tools for Oceanography and Ecosystemic Modeling 143–173 (Wiley, 2016). URL https://onlinelibrary.wiley.com/doi/10.1002/9781119330226.ch5.

[6] Wang, G. Machine learning for inferring animal behavior from location and movement data. Ecological Informatics 49, 69–76 (2019). URL https://linkinghub.elsevier.com/retrieve/pii/S1574954118302036.

[7] Wijeyakulasuriya, D. A., Eisenhauer, E. W., Shaby, B. A. & Hanks, E. M. Machine learning for modeling animal movement. PLOS ONE 15, e0235750 (2020). URL https://journals.plos.org/plosone/article?id=10.1371/journal.pone.0235750.

[8] Potts, J. R. & Börger, L. How to scale up from animal movement decisions to spatiotemporal patterns: An approach via step selection. Journal of Animal Ecology 92, 16–29 (2023). URL https://besjournals.onlinelibrary.wiley.com/doi/10.1111/1365-2656.13832.

[9] Signer, J. et al. Simulating animal space use from fitted integrated Step-Selection Functions (iSSF). Methods in Ecology and Evolution 15, 43–50 (2024). URL https://besjournals.onlinelibrary.wiley.com/doi/10.1111/2041-210X.14263.

[10] Tang, W. & Bennett, D. A. Agent-based modeling of animal movement: a review. Geography Compass 4, 682–700 (2010). URL https://compass.onlinelibrary.wiley.com/doi/10.1111/j.1749-8198.2010.00337.x.

[11] Bergman, C. M., Schaefer, J. A. & Luttich, S. N. Caribou movement as a correlated random walk. Oecologia 123, 364–374 (2000). URL http://link.springer.com/10.1007/s004420051023.

[12] Viswanathan, G., Raposo, E. & Da Luz, M. Lévy flights and superdiffusion in the context of biological encounters and random searches. Physics of Life Reviews 5, 133–150 (2008). URL https://linkinghub.elsevier.com/retrieve/pii/S1571064508000146.

[13] McClintock, B. T. et al. A general discrete-time modeling framework for animal movement using multistate random walks. Ecological Monographs 82, 335–349 (2012). URL https://esajournals.onlinelibrary.wiley.com/doi/10.1890/11-0326.1.

[14] Michelot, T. et al. Estimation and simulation of foraging trips in land-based marine predators. Ecology 98, 1932–1944 (2017). URL https://esajournals.onlinelibrary.wiley.com/doi/10.1002/ecy.1880.

[15] Avgar, T., Potts, J. R., Lewis, M. A. & Boyce, M. S. Integrated step selection analysis: bridging the gap between resource selection and animal movement. Methods in Ecology and Evolution 7, 619–630 (2016). URL https://besjournals.onlinelibrary.wiley.com/doi/10.1111/2041-210X.12528.

[16] Signer, J., Fieberg, J. & Avgar, T. Animal movement tools (amt): R package for managing tracking data and conducting habitat selection analyses. Ecology and Evolution 9, 880–890 (2019). URL https://onlinelibrary.wiley.com/doi/10.1002/ece3.4823.

[17] Nathan, R. et al. A movement ecology paradigm for unifying organismal movement research. Proceedings of the National Academy of Sciences of the United State of America 105, 19052–19059 (2008). URL https://pnas.org/doi/full/10.1073/pnas.0800375105.

[18] Forrest, S. W. et al. Predicting animal movement with deepSSF: a deep learning step selection framework (2025). URL http://biorxiv.org/lookup/doi/10.1101/2025.02.13.638055.

[19] Roy, A., Fablet, R. & Bertrand, S. L. Using generative adversarial networks (GAN) to simulate central-place foraging trajectories. Methods in Ecology and Evolution 13, 1275–1287 (2022). URL https://onlinelibrary.wiley.com/doi/abs/10.1111/2041-210X.13853. xeprint: https://onlinelibrary.wiley.com/doi/pdf/10.1111/2041-210X.13853.

[20] Shaw, A. K. Causes and consequences of individual variation in animal movement. Movement Ecology 8, 12 (2020). URL https://movementecologyjournal.biomedcentral.com/articles/10.1186/s40462-020-0197-x.

[21] Monier, S. A. Social interactions and information use by foraging seabirds. Biological Reviews 99, 1717–1735 (2024). URL https://onlinelibrary.wiley.com/doi/10.1111/brv.13089.

[22] Camphuysen, C. J. Top Predators in Marine Ecosystems: Their Role in Monitoring and Management Conservation Biology (Cambridge University Press, Cambridge, 2006). URL https://www.cambridge.org/core/books/top-predators-in-marine-ecosystems/A4D383B8343181069C9E46A7E783BDFE.

[23] Piatt, I. & Sydeman, W. Seabirds as indicators of marine ecosystems. Marine Ecology Progress Series 352, 199–204 (2007). URL http://www.int-res.com/abstracts/meps/v352/p199-204/.

[24] Arangüena-Proaño, M., Zavalaga, C. & Bugoni, L. Peruvian boobies: unbiased samplers of prey availability and complementary indicators of population size structure of Peruvian anchoveta to fishery landings data. Marine Biology 173, 98 (2026). URL https://link.springer.com/10.1007/s00227-026-04858-x.

[25] Dias, M. P. et al. Threats to seabirds: A global assessment. Biological Conservation 237, 525–537 (2019). URL https://linkinghub.elsevier.com/retrieve/pii/S0006320719307499.

[26] Chen, X., Huang, C., Wang, C. & Chen, L. Trajectory generation: a survey on methods and techniques. GeoInformatica 29, 351–376 (2025). URL https://link.springer.com/10.1007/s10707-025-00545-z.

[27] Bi, J., Cheng, H., Zhang, W., Bao, K. & Wang, P. Artificial intelligence in ship trajectory prediction. Journal of Marine Science and Engineering 12, 769 (2024). URL https://www.mdpi.com/2077-1312/12/5/769. xNumber: 5.

[28] Zhu, Y., Ye, Y., Zhang, S., Zhao, X. & Yu, J. DiffTraj: generating GPS trajectory with diffusion probabilistic model. Advances in Neural Information Processing Systems 36, 65168–65188 (2023). URL https://proceedings.neurips.cc/paper files/paper/2023/hash/cd9b4a28fb9eebe0430c3312a4898a41-Abstract-Conference.html.

[29] Borowiec, M. L. et al. Deep learning as a tool for ecology and evolution. Methods in Ecology and Evolution 13, 1640–1660 (2022). URL https://besjournals.onlinelibrary.wiley.com/doi/10.1111/2041-210X.13901.

[30] Pichler, M. & Hartig, F. Machine learning and deep learning—A review for ecologists. Methods in Ecology and Evolution 14, 994–1016 (2023). URL https://besjournals.onlinelibrary.wiley.com/doi/10.1111/2041-210X.14061.

[31] Tuia, D., Beery, S., Costelloe, B. R., Oliver, R. Y. & Lecomte, N. Towards ‘digital ecology’: advances in integrating artificial intelligence from data generation to ecological insight. Methods in Ecology and Evolution 17, 222–227 (2026). URL https://besjournals.onlinelibrary.wiley.com/doi/10.1111/2041-210x.70243.

[32] Schwartz, R., Dodge, J., Smith, N. A. & Etzioni, O. Green AI. Communications of the ACM 63, 54–63 (2020). URL https://dl.acm.org/doi/10.1145/3381831.

[33] Joo, R. et al. Recent trends in movement ecology of animals and human mobility. Movement Ecology 10, 26 (2022). URL https://movementecologyjournal.biomedcentral.com/articles/10.1186/s40462-022-00322-9.

[34] Ding, N. et al. Parameter-efficient fine-tuning of large-scale pre-trained language models. Nature Machine Intelligence 5, 220–235 (2023). URL https://www.nature.com/articles/s42256-023-00626-4.

[35] Zhuang, F. et al. A comprehensive survey on transfer learning. Proceedings of the IEEE 109, 43–76 (2021). URL https://ieeexplore.ieee.org/document/9134370/.

[36] Devlin, J., Chang, M.-W., Lee, K. & Toutanova, K. BERT: pre-training of deep bidirectional transformers for language understanding. Proceedings of the 2019 Conference of the North American Chapter of the Association for Computational Linguistics: Human Language Technologies, Volume 1 (Long and Short Papers) 4171–4186 (2019).

[37] Yuan, L. et al. Florence: a new foundation model for computer vision (2021). URL http://arxiv.org/abs/2111.11432. ArXiv:2111.11432 [cs.CV].

[38] Sohl-Dickstein, J., Weiss, E. A., Maheswaranathan, N. & Ganguli, S. Deep unsupervised learning using nonequilibrium thermodynamics. Proceedings of the 32nd International Conference on Machine Learning (2015). URL https://proceedings.mlr.press/v37/sohl-dickstein15.html.

[39] Ho, J., Jain, A. & Abbeel, P. Denoising diffusion probabilistic models (2020). URL http://arxiv.org/abs/2006.11239. ArXiv:2006.11239 [cs].

[40] Yang, Y. et al. A survey on diffusion models for time series and spatio-temporal data (2024). URL http://arxiv.org/abs/2404.18886. ArXiv:2404.18886 [cs].

[41] Yang, L. et al. Diffusion models: a comprehensive survey of methods and applications. ACM Computing Surveys 56, 1–39 (2024). URL https://dl.acm.org/doi/10.1145/3626235.

[42] Bost, C. et al. The importance of oceanographic fronts to marine birds and mammals of the southern oceans. Journal of Marine Systems 78, 363–376 (2009). URL https://linkinghub.elsevier.com/retrieve/pii/S0924796309000724.

[43] Embling, C. B. et al. Investigating fine-scale spatio-temporal predator–prey patterns in dynamic marine ecosystems: a functional data analysis approach. Journal of Applied Ecology 49, 481–492 (2012). URL https://besjournals.onlinelibrary.wiley.com/doi/10.1111/j.1365-2664.2012.02114.x.

[44] Scales, K. L. et al. Mesoscale fronts as foraging habitats: composite front mapping reveals oceanographic drivers of habitat use for a pelagic seabird. Journal of The Royal Society Interface 11, 20140679 (2014). URL https://royalsocietypublishing.org/doi/10.1098/rsif.2014.0679.

[45] O’Hanlon, N. J. et al. Challenges in quantifying the responses of Black-legged Kittiwakes Rissa tridactyla to habitat variables and local stressors due to individual variation. Bird Study 71, 48–64 (2024). URL https://www.tandfonline.com/doi/full/10.1080/00063657.2024.2305169.

[46] Keys, D., Pistorius, P., Tremblay, Y. & Thiebault, A. Both wind and flying with conspecifics influence the flight dynamics of a seabird. Marine Ecology Progress Series 723, 135–149 (2023). URL https://www.int-res.com/abstracts/meps/v723/p135-149/.

[47] Dunn, R. E. et al. Commuting in crosswinds and foraging in fast winds: the foraging ecology of a flying fish specialist. Proceedings of the Royal Society B: Biological Sciences 292, 20250774 (2025). URL https://royalsocietypublishing.org/doi/10.1098/rspb.2025.0774.

[48] Niven, H. I., Jeglinski, J. W. E., Aarts, G., Wakefield, E. D. & Matthiopoulos, J. Towards biologically realistic estimates of home range and spatial exposure for colonial animals. Methods in Ecology and Evolution 2041–210X.70019 (2025). URL https://besjournals.onlinelibrary.wiley.com/doi/10.1111/2041-210X.70019.

[49] Zucchini, W., MacDonald, I. & Langrock, R. Hidden Markov Models for Time Series: An Introduction Using R Chapman and Hall/CRC Monographs on Statistics and Applied Probability Series (CRC Press, Taylor & Francis Group, 2016). URL https://books.google.fr/books?id=LT3RjwEACAAJ.

[50] Vivekananthan, S. Comparative analysis of generative models: enhancing image synthesis with VAEs, GANs, and Stable Diffusion (2024). URL http://arxiv.org/abs/2408.08751. ArXiv:2408.08751 [cs.CV].

[51] Wang, H. Comparative analysis of GANs and Diffusion models in image generation. Highlights in Science, Engineering and Technology 120, 59–66 (2024). URL https://drpress.org/ojs/index.php/HSET/article/view/28780.

[52] McClintock, B. T. & Michelot, T. momentuHMM: R package for generalized hidden Markov models of animal movement. Methods in Ecology and Evolution 9, 1518–1530 (2018). URL https://besjournals.onlinelibrary.wiley.com/doi/10.1111/2041-210X.12995.

[53] Langrock, R. et al. Flexible and practical modeling of animal telemetry data: hidden Markov models and extensions. Ecology 93, 2336–2342 (2012). URL https://esajournals.onlinelibrary.wiley.com/doi/10.1890/11-2241.1.

[54] Thurfjell, H., Ciuti, S. & Boyce, M. S. Applications of step-selection functions in ecology and conservation. Movement Ecology 2, 4 (2014). URL https://movementecologyjournal.biomedcentral.com/articles/10.1186/2051-3933-2-4.

[55] Li, H., Jiao, H. & Yang, Z. Ship trajectory prediction based on machine learning and deep learning: A systematic review and methods analysis. Engineering Applications of Artificial Intelligence 126, 107062 (2023). URL https://linkinghub.elsevier.com/retrieve/pii/S0952197623012460.

[56] Pollock, C. J. et al. Movement simulations reveal both memory and social information drive individual foraging site fidelity in gannets. Movement Ecology (2026). URL 10.1186/s40462-026-00656-8.

[57] Schreiber, E. A. (ed.) Biology of marine birds CRC marine biology series (CRC Press, Boca Raton, Fla., 2002).

[58] Bolnick, D. et al. The ecology of individuals: incidence and implications of individual specialization. The American Naturalist 161, 1–28 (2003). URL https://www.journals.uchicago.edu/doi/10.1086/343878.

[59] Weimerskirch, H., Bertrand, S., Silva, J., Marques, J. C. & Goya, E. Use of social information in seabirds: compass rafts indicate the heading of food patches. PLoS ONE 5, e9928 (2010). URL 10.1371/journal.pone.0009928.

[60] Hindell, M. A. et al. Tracking of marine predators to protect Southern Ocean ecosystems. Nature 580, 87–92 (2020). URL https://www.nature.com/articles/s41586-020-2126-y.

[61] Wakefield, E., Phillips, R. & Matthiopoulos, J. Quantifying habitat use and preferences of pelagic seabirds using individual movement data: a review. Marine Ecology Progress Series 391, 165–182 (2009). URL http://www.int-res.com/abstracts/meps/v391/p165-182/.

[62] Tremblay, Y. et al. Analytical approaches to investigating seabird–environment interactions: a review. Marine Ecology Progress Series 391, 153–163 (2009). URL http://www.int-res.com/abstracts/meps/v391/p153-163/.

[63] Critchley, E. J., Grecian, W. J., Kane, A., Jessopp, M. J. & Quinn, J. L. Marine protected areas show low overlap with projected distributions of seabird populations in Britain and Ireland. Biological Conservation 224, 309–317 (2018). URL https://linkinghub.elsevier.com/retrieve/pii/S0006320717317469.

[64] Garthe, S. et al. Wind farms in proximity to marine protected areas put conservation targets at risk. Journal for Nature Conservation 84, 126805 (2025). URL https://linkinghub.elsevier.com/retrieve/pii/S1617138124002541.

[65] Dierschke, V., Furness, R. W. & Garthe, S. Seabirds and offshore wind farms in European waters: Avoidance and attraction. Biological Conservation 202, 59–68 (2016). URL https://linkinghub.elsevier.com/retrieve/pii/S0006320716303196.

[66] Oppel, S. et al. Spatial scales of marine conservation management for breeding seabirds. Marine Policy 98, 37–46 (2018). URL https://linkinghub.elsevier.com/retrieve/pii/S0308597X18302422.

[67] Wakefield, E. D. et al. Long-term individual foraging site fidelity—why some gannets don’t change their spots. Ecology 96, 3058–3074 (2015). URL https://esajournals.onlinelibrary.wiley.com/doi/10.1890/14-1300.1.

[68] Michelot, T., Langrock, R. & Patterson, T. A. moveHMM: an R package for the statistical modelling of animal movement data using hidden Markov models. Methods in Ecology and Evolution 7, 1308–1315 (2016). URL https://besjournals.onlinelibrary.wiley.com/doi/10.1111/2041-210X.12578.

