## Supplementary material for "From vehicles to wildlife: transferable deep learning for trajectory generation"

### A Durations of training and simulation

**Table 1**: Training duration of generative trajectory models across dataset subsets. The diffusion model is trained once on the full pooled dataset; GAN, VAE, HMM, and iSSF are trained independently per subset. HMM and iSSF are trained only on regularized GPS points, while diffusion, GAN and VAE are trained on padded trajectories, all composed of 200 points. ep. = epochs; min = minutes; s = seconds. HMM timing excludes initialization search; iSSF timing is dominated by variable extraction from control steps.

| | **Brazil, Abrolhos** | | **Brazil, Fernando de Noronha** | | | **Brazil, Santana** | **Brazil, São Pedro e São Paulo** | **Peru, Guañape** | **Peru, Pescadores** | | |
|---|---|---|---|---|---|---|---|---|---|---|---|
| | Masked booby | Brown booby | Masked booby | Brown booby | Red-footed booby | Brown booby | Brown booby | Peruvian booby | Guanay cormorant | Peruvian booby | **Total** |
| Nb. trajectories | 98 | 38 | 187 | 14 | 48 | 145 | 447 | 47 | 194 | 412 | 1 630 |
| Nb. GPS points | 7 497 | 1 899 | 11 599 | 971 | 6 672 | 3 761 | 8 316 | 1 739 | 5 968 | 7 633 | 56 055 |
| *Trained on full dataset* | | | | | | | | | | | |
| Diffusion fine-tuning (200 ep.) | | | | | | 35 min | | | | | 35 min |
| Diffusion from scratch (3 000 ep.) | | | | | | 7 h | | | | | 7 h |
| *Trained per subset* | | | | | | | | | | | |
| GAN (5 000 ep.) | 2 min | 1 min | 3 min 10 s | 30 s | 1 min 20 s | 2 min 50 s | 6 min 50 s | 1 min 20 s | 3 min 20 s | 6 min 20 s | 29 min |
| VAE (1 000 ep.) | 1 min 30 s | 50 s | 2 min 20 s | 30 s | 1 min | 1 min 50 s | 6 min | 1 min | 2 min 30 s | 5 min 30 s | 23 min |
| HMM | 6 min | 1 min 40 s | 15 min | 1 min | 6 min | 3 min | 5 min | 1 min 20 s | 5 min | 7 min | 51 min |
| iSSF | 1 min | 20 s | 1 min 30 s | 10 s | 50 s | 30 s | 1 min 10 s | 20 s | 40 s | 1 min | 7 min 30 s |

**Table 2**: Generation and post-processing duration per method across dataset subsets, for $10 \times n_{\text{traj,real}}$ generated trajectories. Diffusion generation time is shared across all subsets. GAN generation is near-instant; only post-processing is timed. VAE generation requires negligible time. HMM post-processing retains only non-looping, on-earth trajectories. min = minutes; s = seconds; h = hours.

| | **Brazil, Abrolhos** | | **Brazil, Fernando de Noronha** | | | **Brazil, Santana** | **Brazil, São Pedro e São Paulo** | **Peru, Guañape** | **Peru, Pescadores** | | |
|---|---|---|---|---|---|---|---|---|---|---|---|
| | Masked booby | Brown booby | Masked booby | Brown booby | Red-footed booby | Brown booby | Brown booby | Peruvian booby | Guanay cormorant | Peruvian booby | **Total** |
| Nb. trajectories | 98 | 38 | 187 | 14 | 48 | 145 | 447 | 47 | 194 | 412 | 1 630 |
| Nb. GPS points | 7 497 | 1 899 | 11 599 | 971 | 6 672 | 3 761 | 8 316 | 1 739 | 5 968 | 7 633 | 56 055 |
| *Trained on full dataset* | | | | | | | | | | | |
| Diffusion fine-tuning (200 ep.) | | | | | | 30 min | | | | | 30 min |
| Diffusion from scratch (3 000 ep.) | | | | | | 30 min | | | | | 30 min |
| *Trained per subset* | | | | | | | | | | | |
| GAN (post-processing only) | 15 s | 5 s | 30 s | 0 s | 10 s | 20 s | 2 min 30 s | 10 s | 30 s | 2 min | 6 min 30 s |
| VAE | | | | | | 0 s | | | | | 0 s |
| HMM | 17 min | 6 min | 33 min | 3 min | 10 min | 23 min | 1 h 10 min | 7 min | 33 min | 1 h 15 min | 4 h 37 min |
| iSSF | 33 min | 6 min | 41 min | 3 min 30 s | 11 min 30 s | 18 min | 40 min | 6 min 30 s | 20 min | 1 h | 4 h |

#### B Diffusion VS State-of-the-art

We show here the results from all the subsets. The boxplots were obtained by simulating 10 times the number of real trajectories, and by evaluating each of the 10 simulated sets obtained separately. This aims at taking into account the variability associated with the simulation process, especially for small subsets.

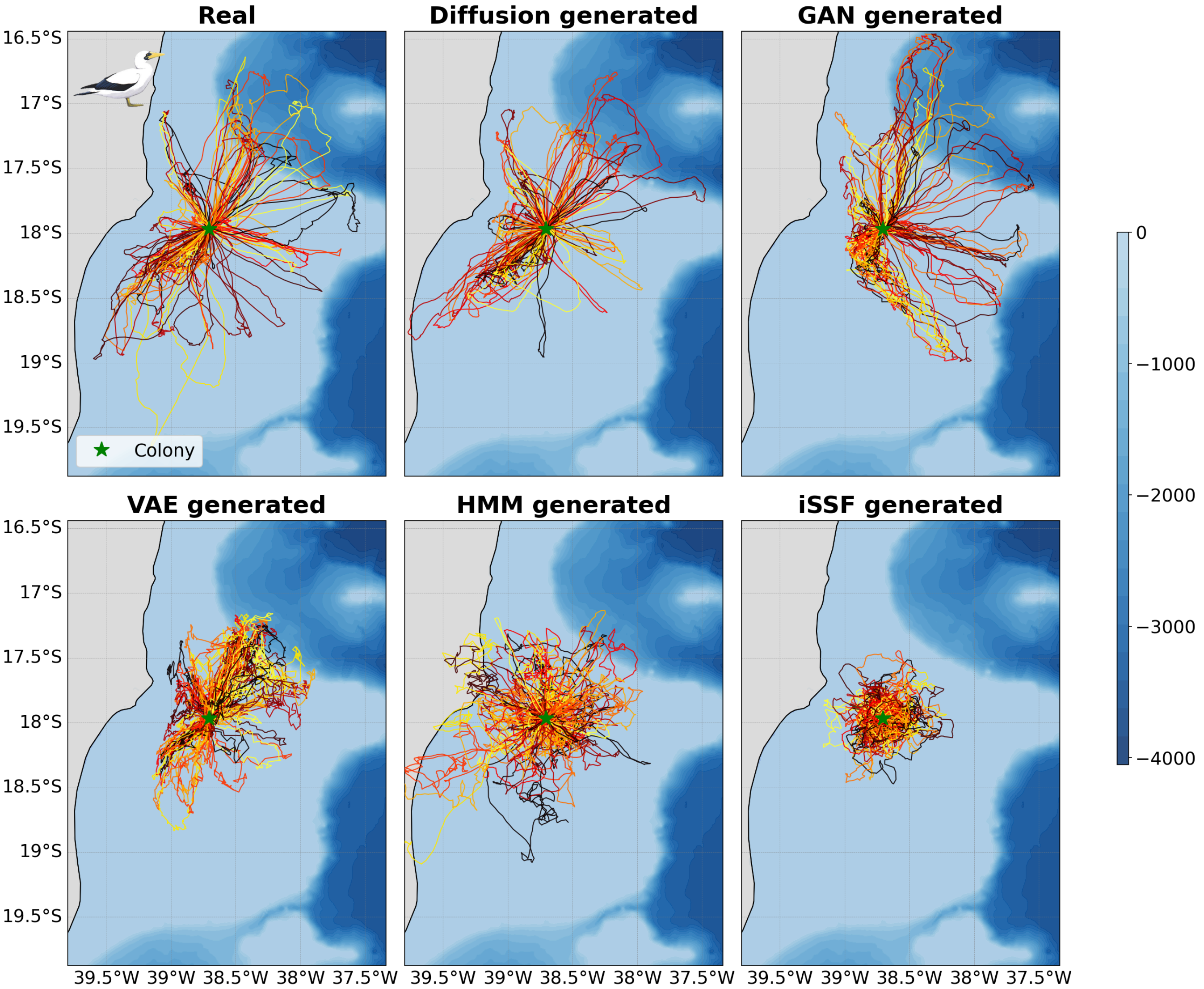


**Fig. 1**: **Real and model-generated trajectories for Masked Boobies (*Sula dactylatra*) at Abrolhos Archipelago, Brazil.** Each panel shows 98 trajectories. A trajectory is an out-and-back trip from the colony. Colors represent unique trips and are randomized for visual clarity.

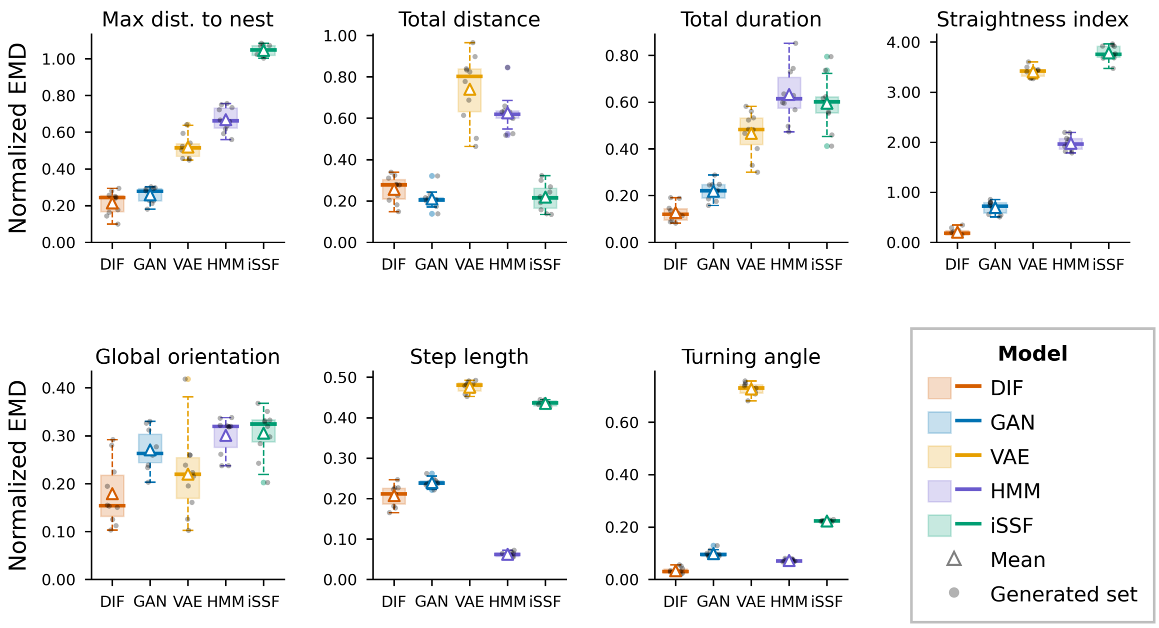


(a) Per-metrics errors.

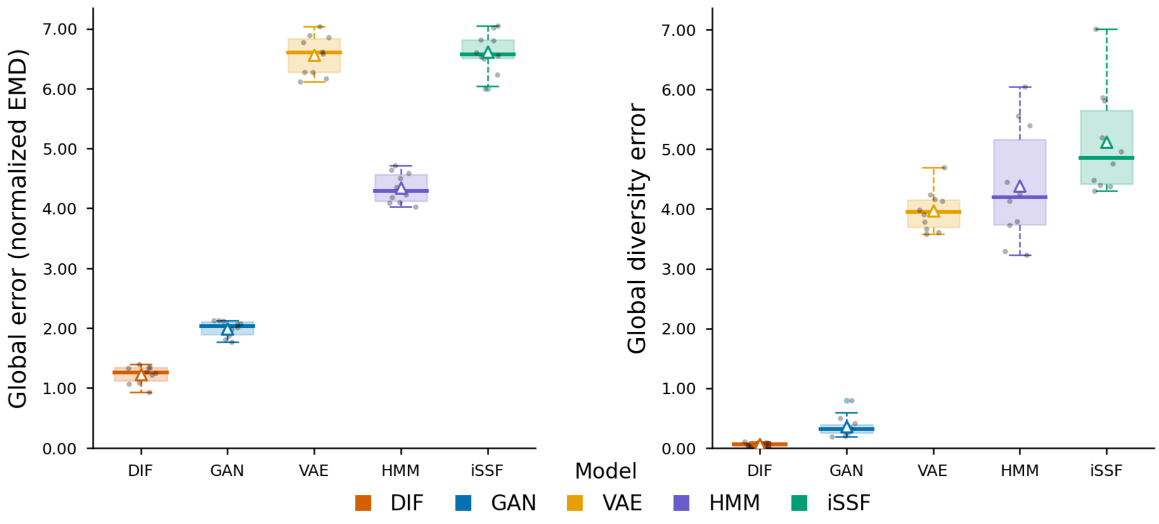


(b) Global scores.

**Fig. 2**: **Model comparison for Masked Boobies (*Sula dactylatra*) at Abrolhos Archipelago, Brazil ($n = 98$).** Panels show a) per-metric errors and b) global scores, computed as the Earth Mover's Distance between real and generated distributions across 10 independently generated sets per model.

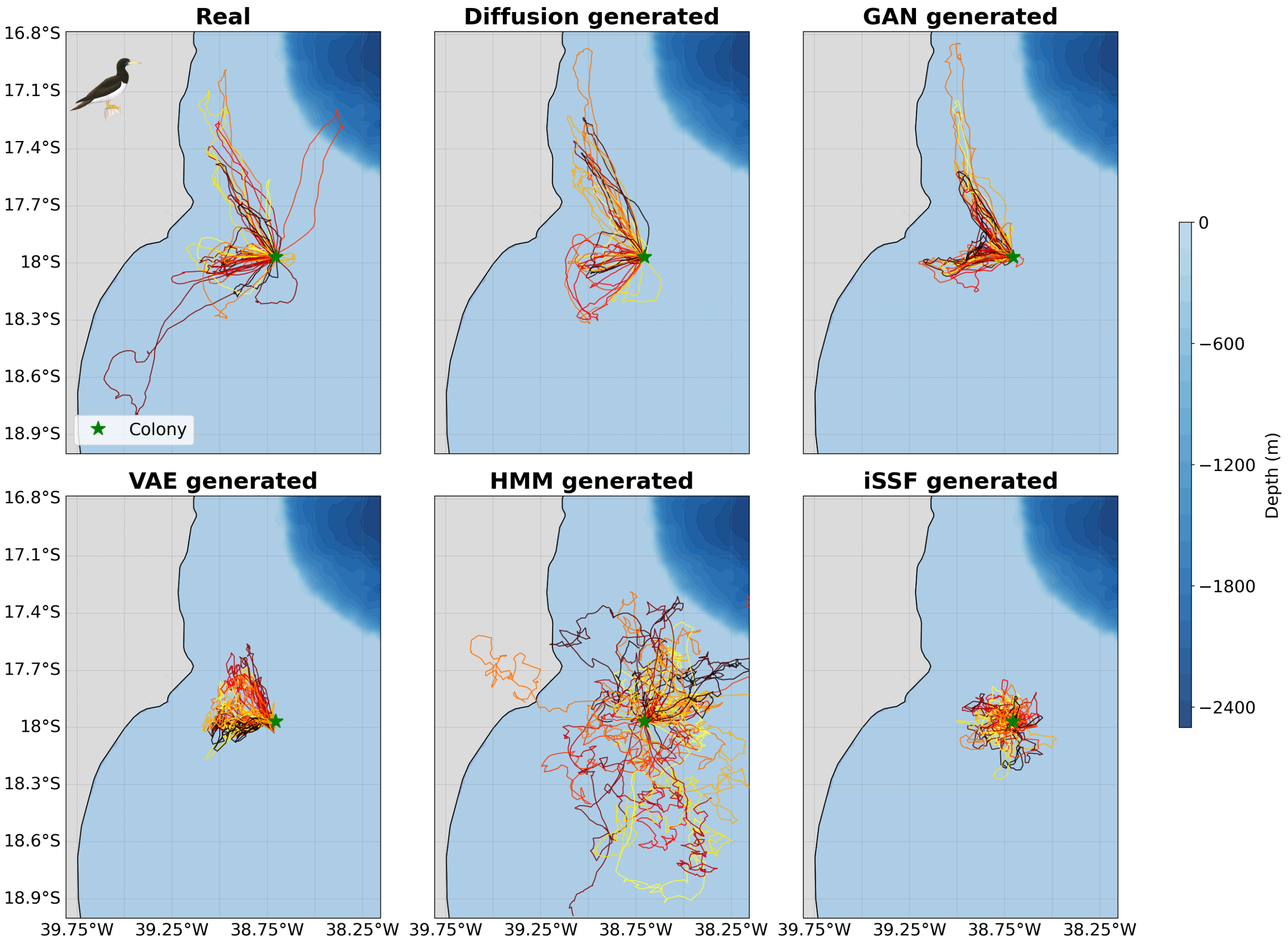


**Fig. 3**: **Real and model-generated trajectories for Brown Boobies (*Sula leucogaster*) at Abrolhos Archipelago, Brazil.** Each panel shows 38 trajectories. A trajectory is an out-and-back trip from the colony. Colors represent unique trips and are randomized for visual clarity.

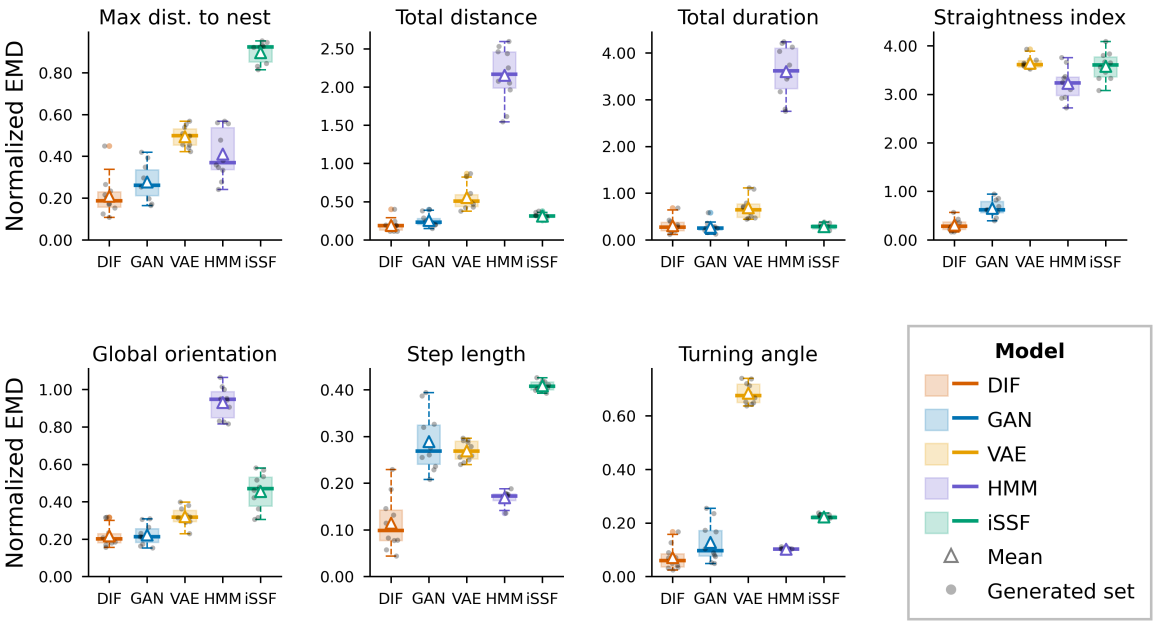


(a) Per-metrics errors.

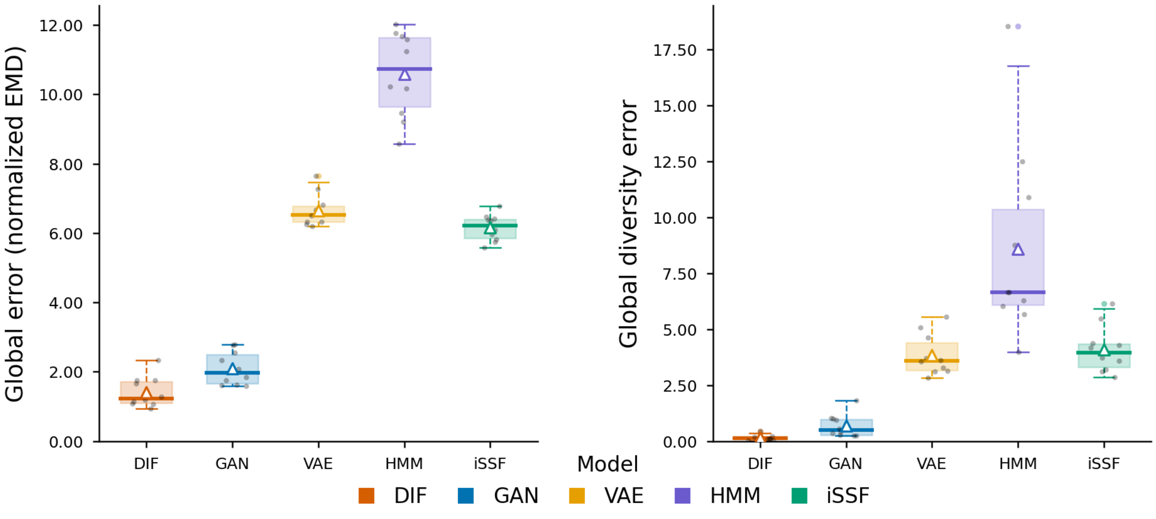


(b) Global scores.

**Fig. 4**: **Model comparison for Brown Boobies (*Sula leucogaster*) at Abrolhos Archipelago, Brazil ($n = 38$).** Panels show a) per-metric errors and b) global scores, computed as the Earth Mover's Distance between real and generated distributions across 10 independently generated sets per model.

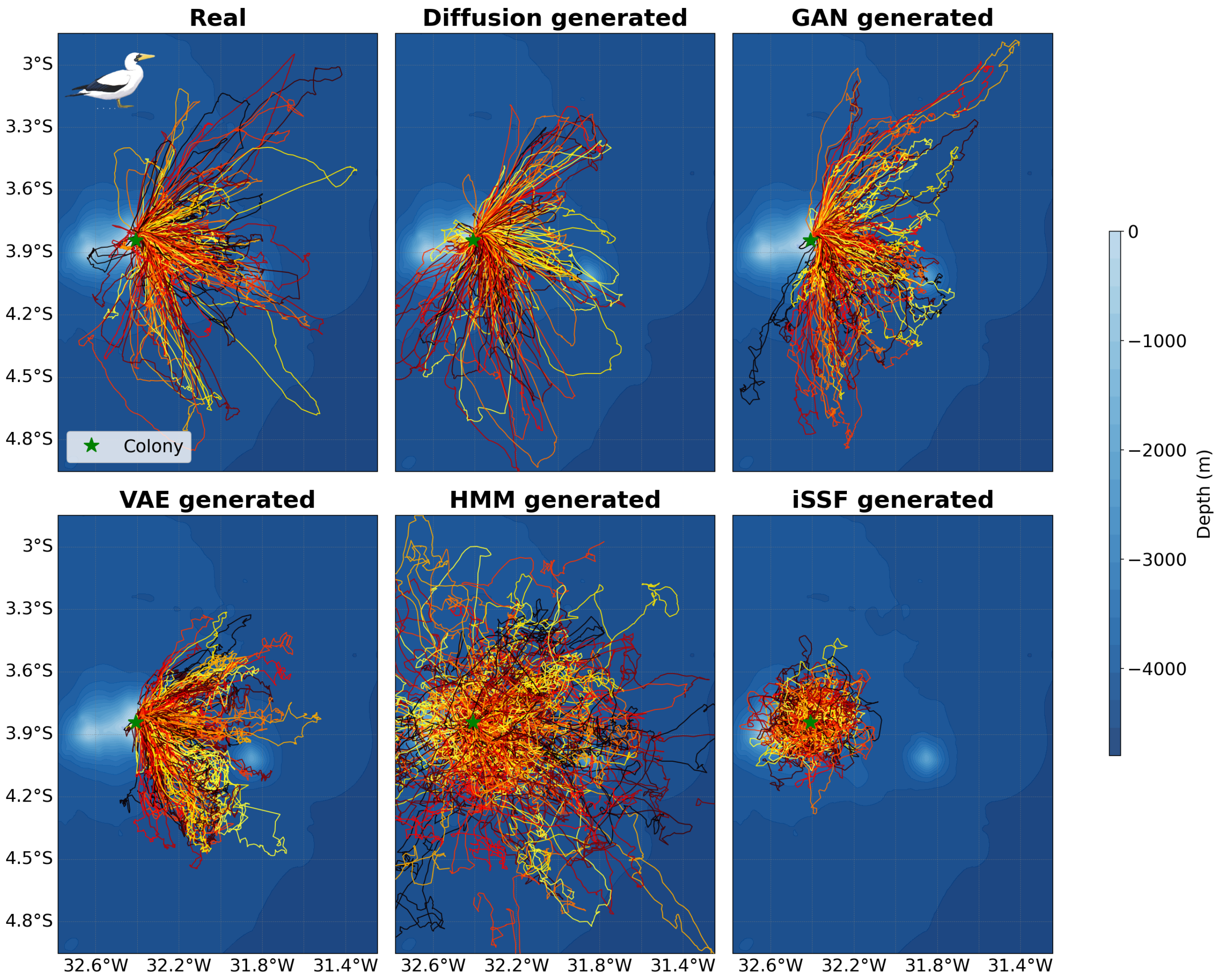


**Fig. 5**: **Real and model-generated trajectories for Masked Boobies (*Sula dactylatra*) at Fernando de Noronha, Brazil.** Each panel shows 187 trajectories. A trajectory is an out-and-back trip from the colony. Colors represent unique trips and are randomized for visual clarity.

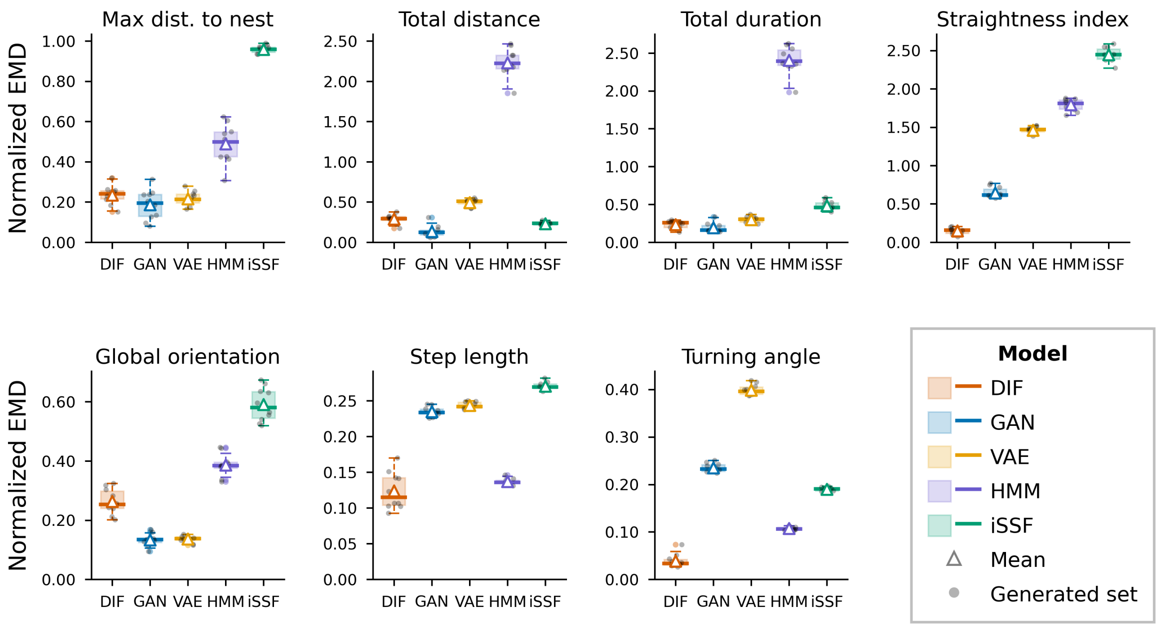


(a) Per-metrics errors.

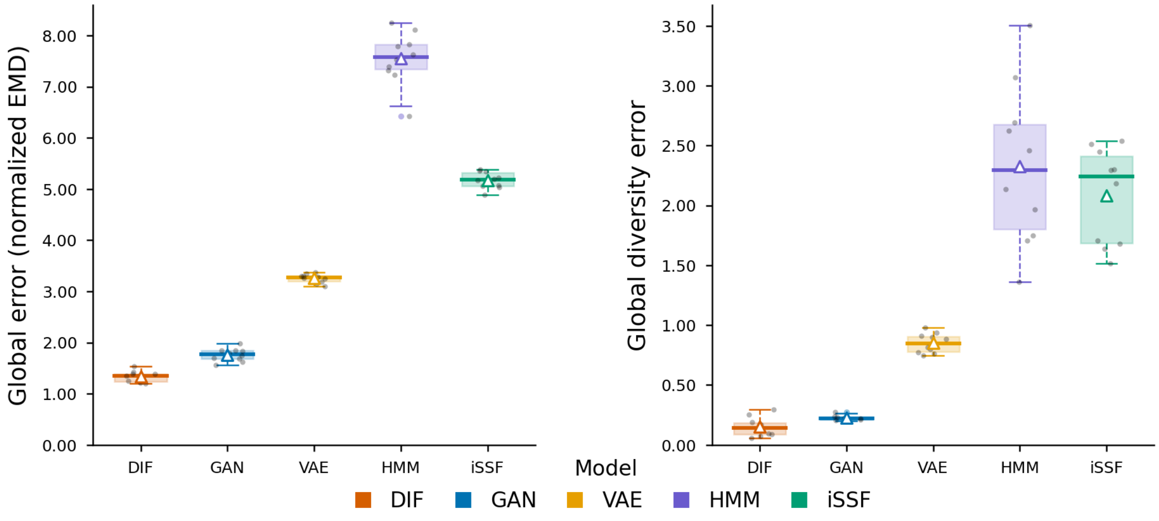


(b) Global scores.

**Fig. 6**: **Model comparison for Masked Boobies (*Sula dactylatra*) at Fernando de Noronha, Brazil ($n$ = 187).** Panels show a) per-metric errors and b) global scores, computed as the Earth Mover's Distance between real and generated distributions across 10 independently generated sets per model.

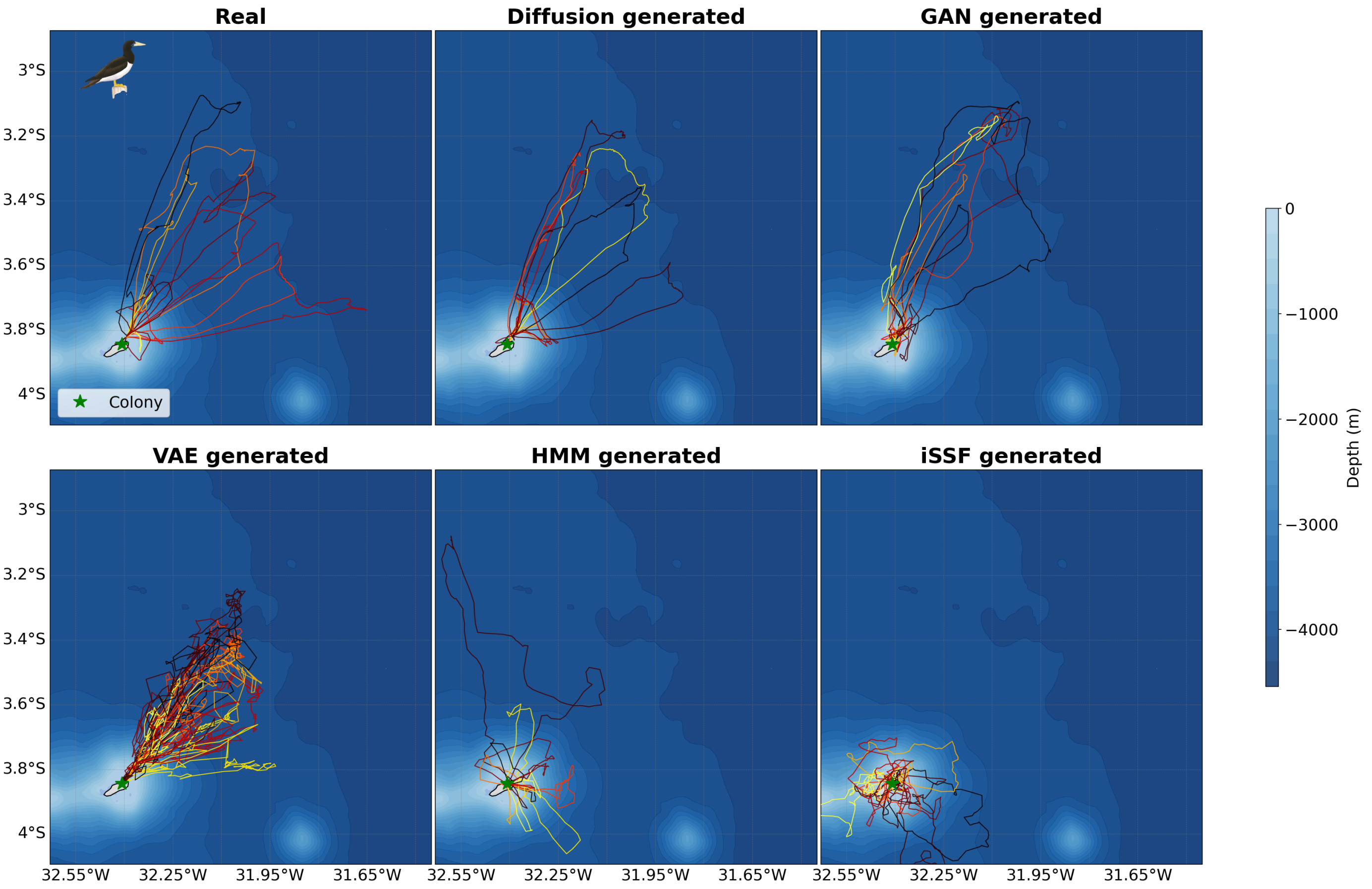


**Fig. 7**: **Real and model-generated trajectories for Brown Boobies (*Sula leucogaster*) at Fernando de Noronha, Brazil.** Each panel shows 14 trajectories. A trajectory is an out-and-back trip from the colony. Colors represent unique trips and are randomized for visual clarity.

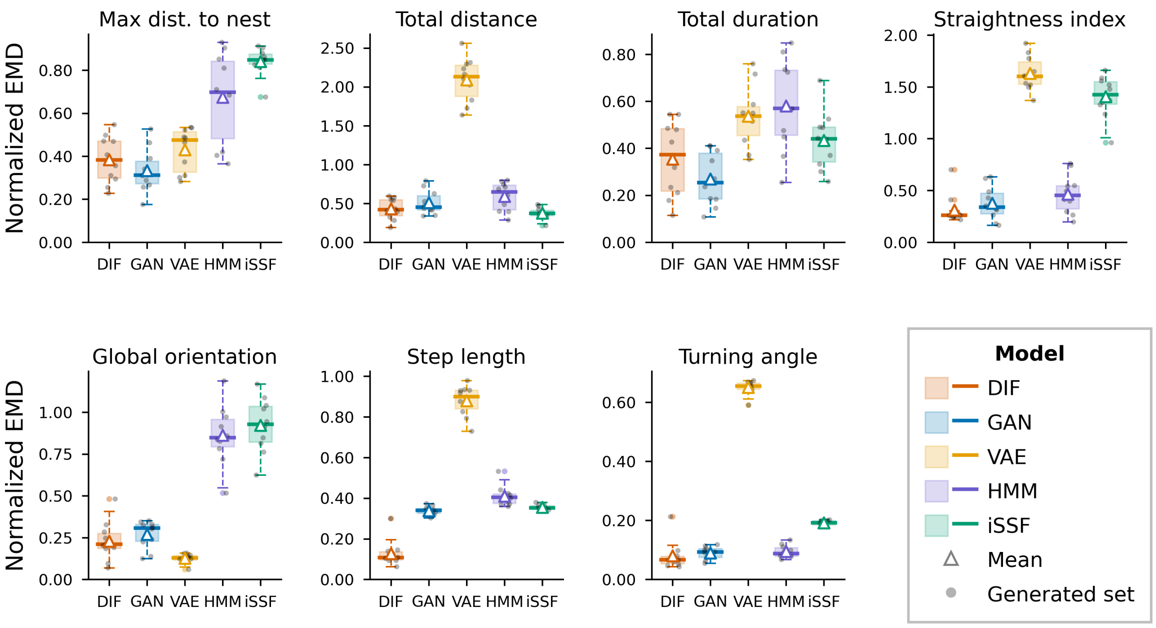


(a) Per-metrics errors.

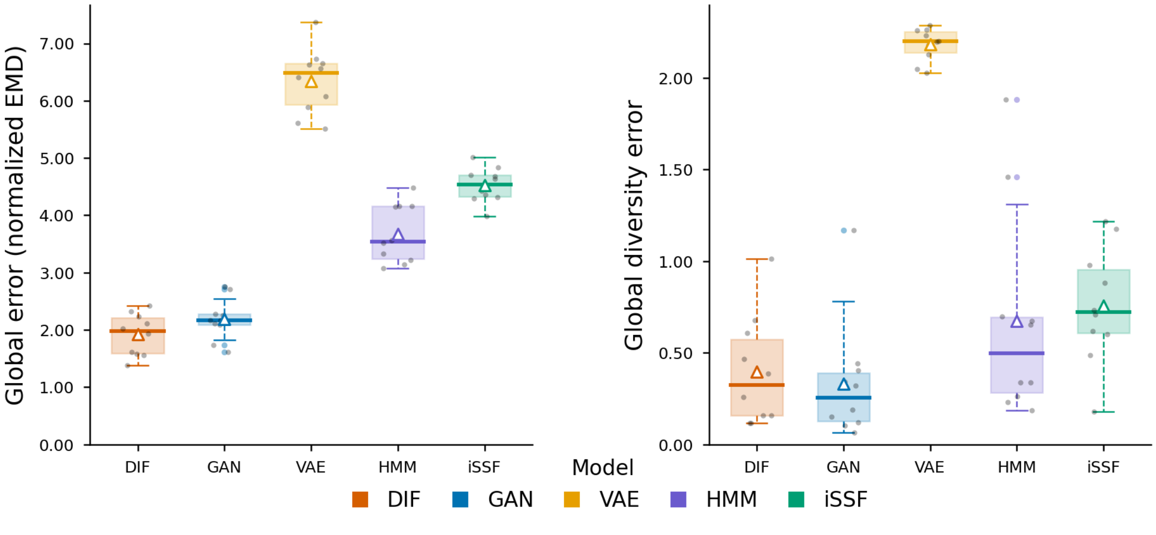


(b) Global scores.

**Fig. 8**: **Model comparison for Brown Boobies (*Sula leucogaster*) at Fernando de Noronha, Brazil ($n = 14$).** Panels show a) per-metric errors and b) global scores, computed as the Earth Mover's Distance between real and generated distributions across 10 independently generated sets per model.

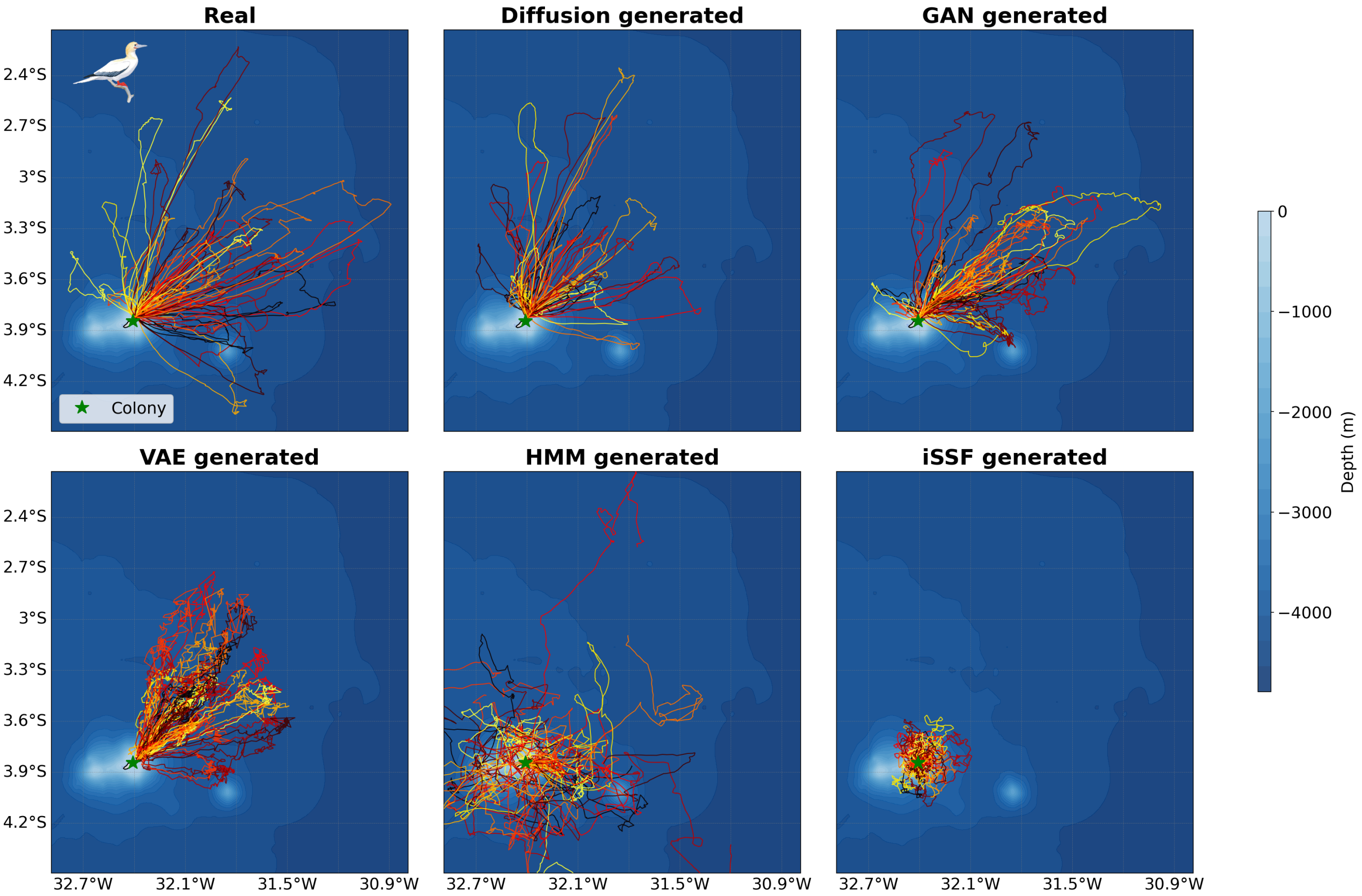


**Fig. 9**: **Real and model-generated trajectories for Red-footed Boobies (*Sula sula*) at Fernando de Noronha, Brazil.** Each panel shows 48 trajectories. A trajectory is an out-and-back trip from the colony. Colors represent unique trips and are randomized for visual clarity.

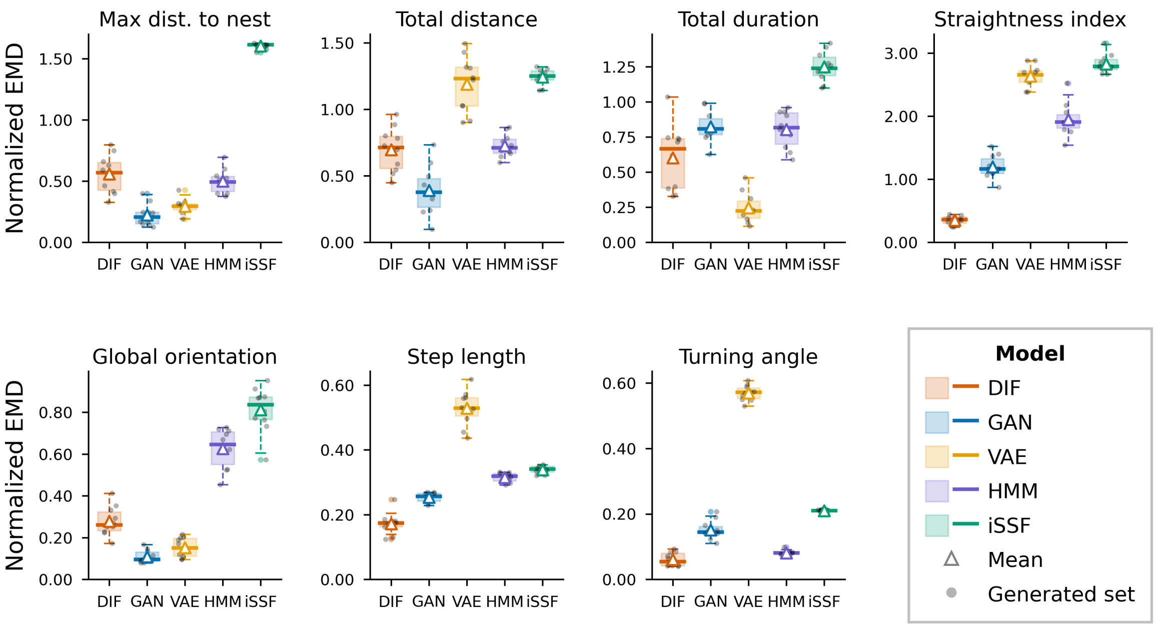


(a) Per-metrics errors.

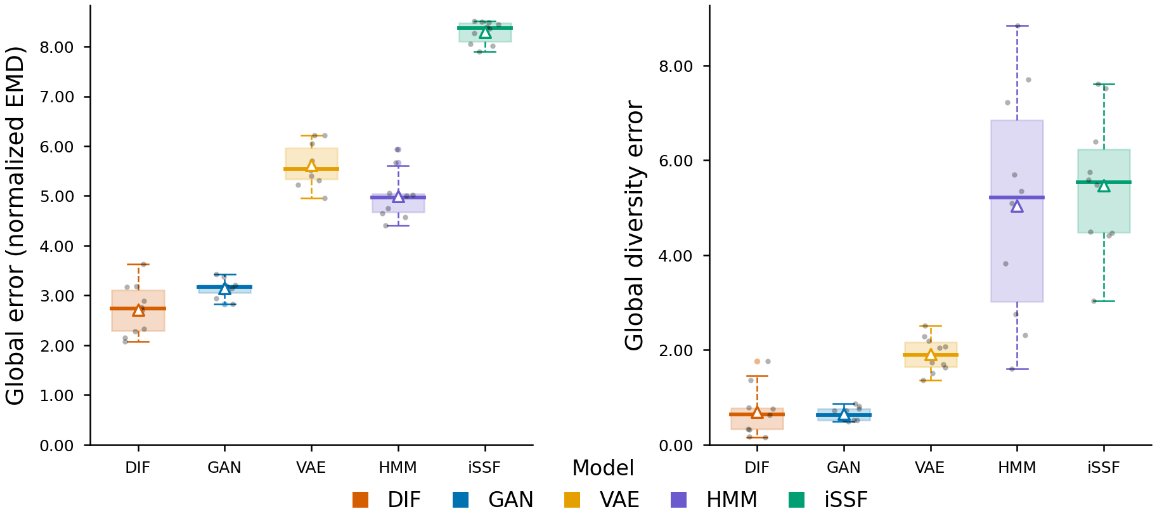


(b) Global scores.

**Fig. 10**: **Model comparison for Red-footed Boobies (*Sula sula*) at Fernando de Noronha, Brazil ($n$ = 48).** Panels show a) per-metric errors and b) global scores, computed as the Earth Mover's Distance between real and generated distributions across 10 independently generated sets per model.

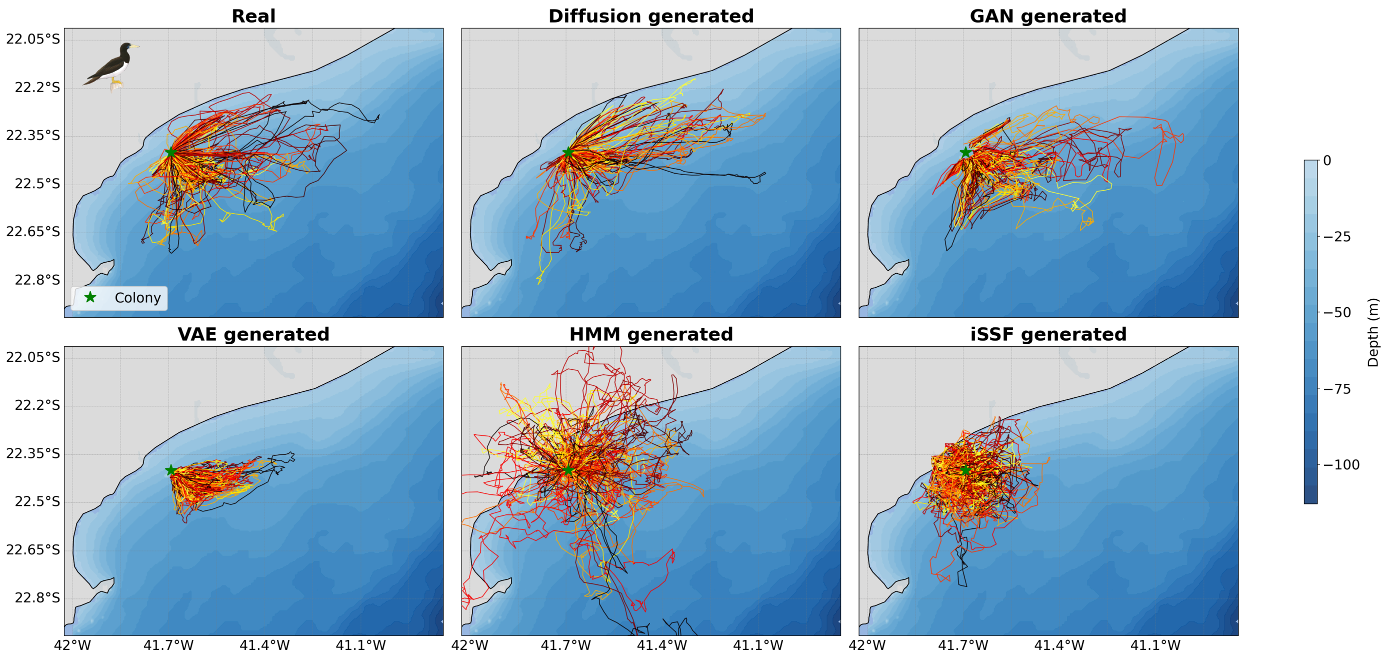


**Fig. 11**: **Real and model-generated trajectories for Brown Boobies (*Sula leucogaster*) at Santana Archipelago, Brazil.** Each panel shows 145 trajectories. A trajectory is an out-and-back trip from the colony. Colors represent unique trips and are randomized for visual clarity.

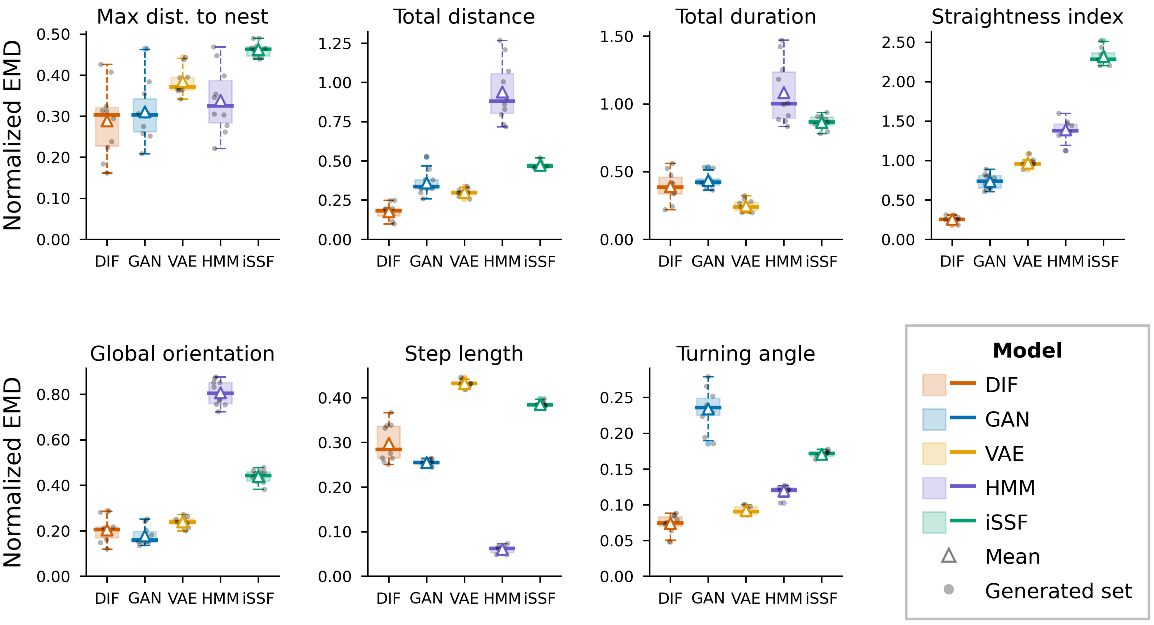


(a) Per-metrics errors.

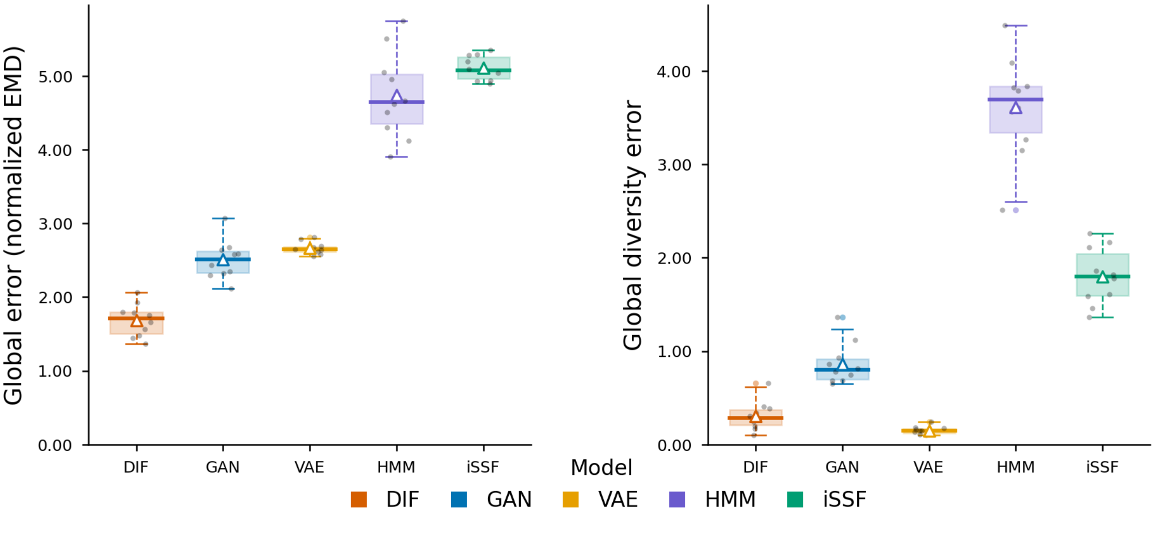


(b) Global scores.

**Fig. 12**: **Model comparison for Brown Boobies (*Sula leucogaster*) at Santana Archipelago, Brazil ($n = 145$).** Panels show a) per-metric errors and b) global scores, computed as the Earth Mover's Distance between real and generated distributions across 10 independently generated sets per model.

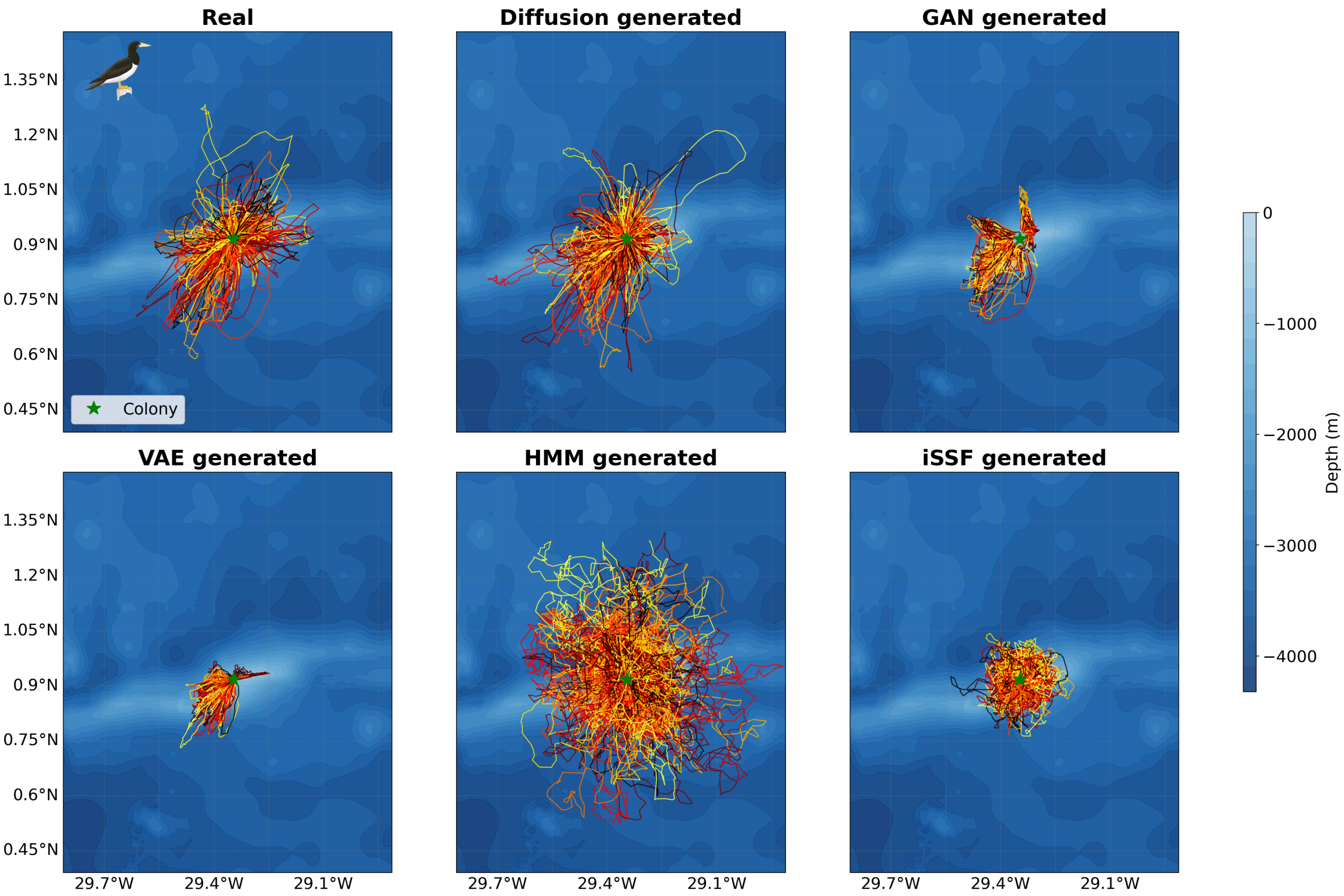


**Fig. 13**: **Real and model-generated trajectories for Brown Boobies (*Sula leucogaster*) at São Pedro e São Paulo Archipelago, Brazil.** Each panel shows 447 trajectories. A trajectory is an out-and-back trip from the colony. Colors represent unique trips and are randomized for visual clarity.

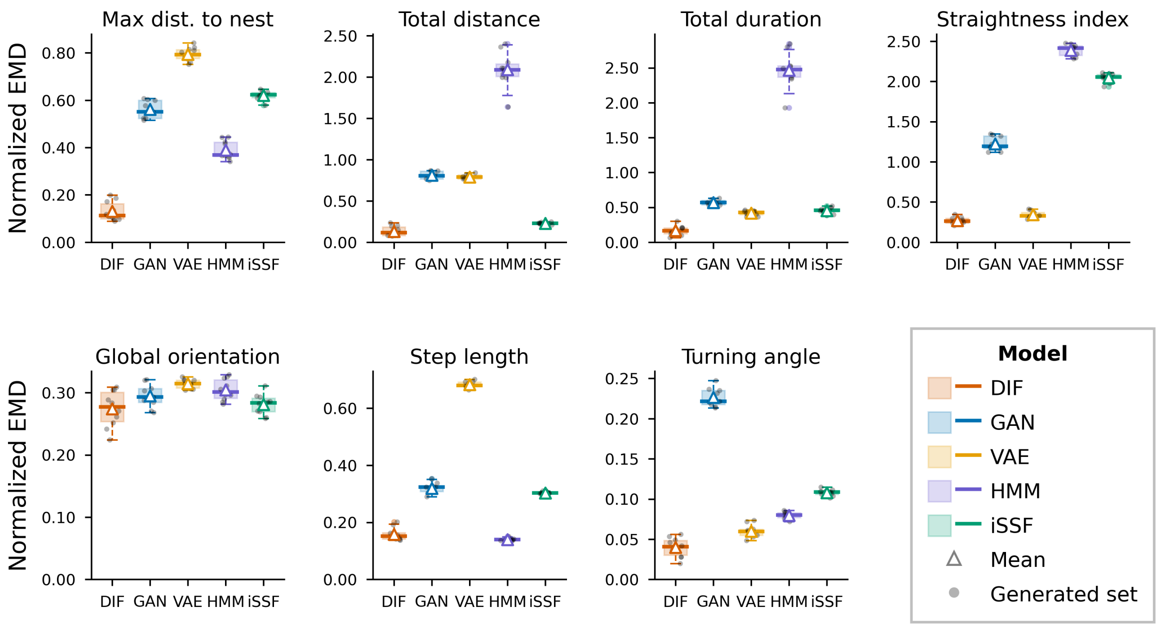


(a) Per-metrics errors.

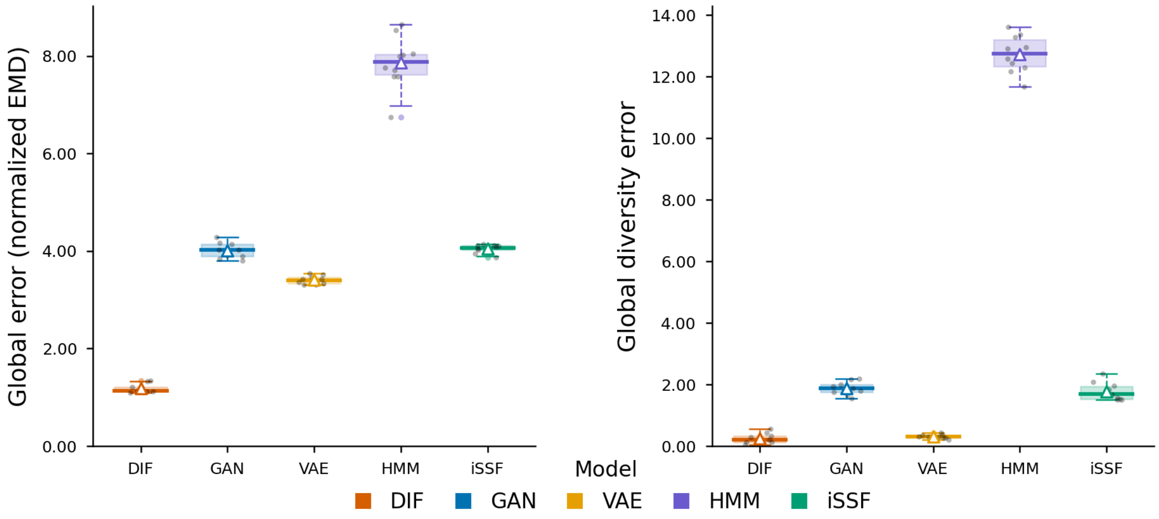


(b) Global scores.

**Fig. 14**: **Model comparison for Brown Boobies (*Sula leucogaster*) at São Pedro e São Paulo Archipelago, Brazil ($n$ = 447).** Panels show a) per-metric errors and b) global scores, computed as the Earth Mover's Distance between real and generated distributions across 10 independently generated sets per model.

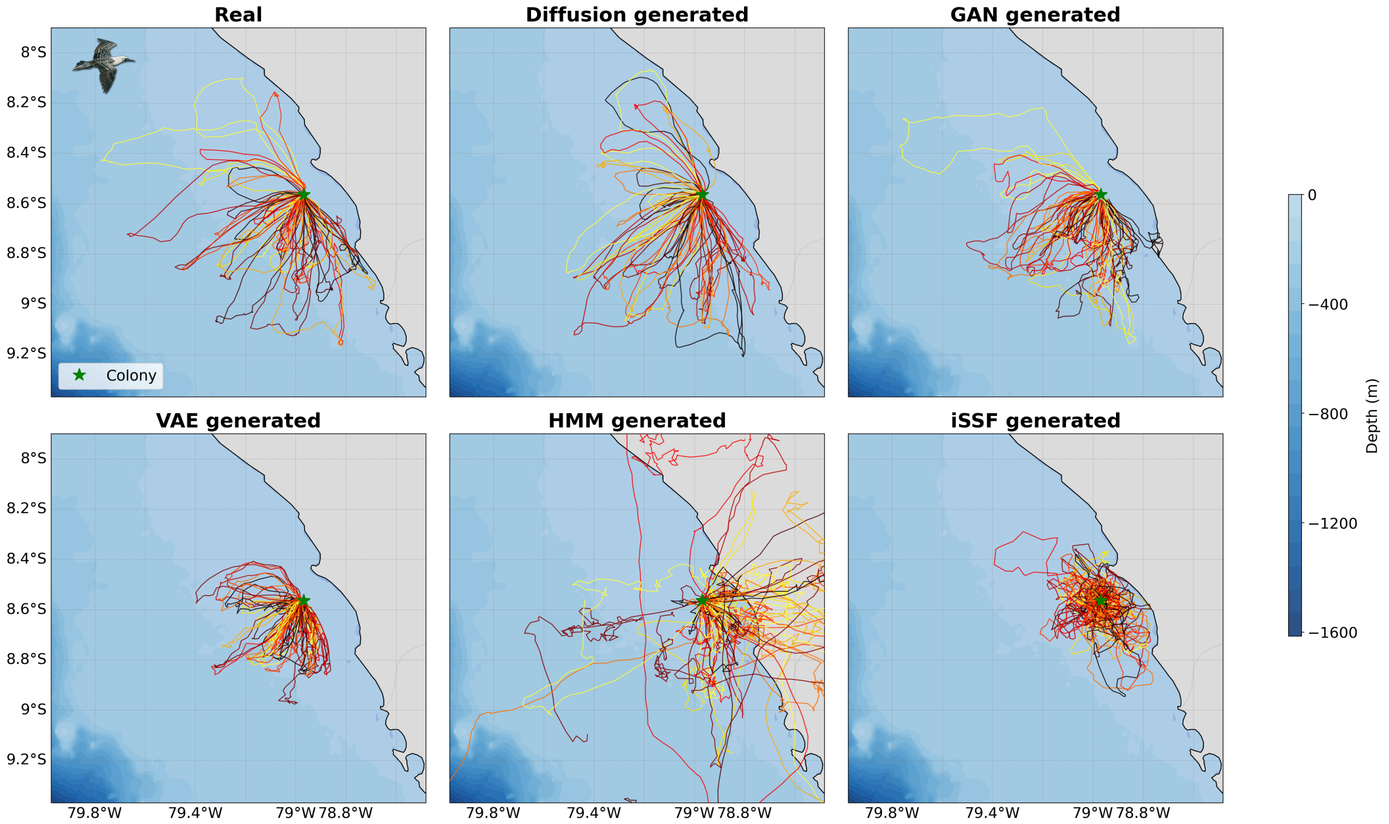


**Fig. 15**: **Real and model-generated trajectories for Peruvian Boobies (*Sula variegata*) at Guañape, Peru.** Each panel shows 47 trajectories. A trajectory is an out-and-back trip from the colony. Colors represent unique trips and are randomized for visual clarity.

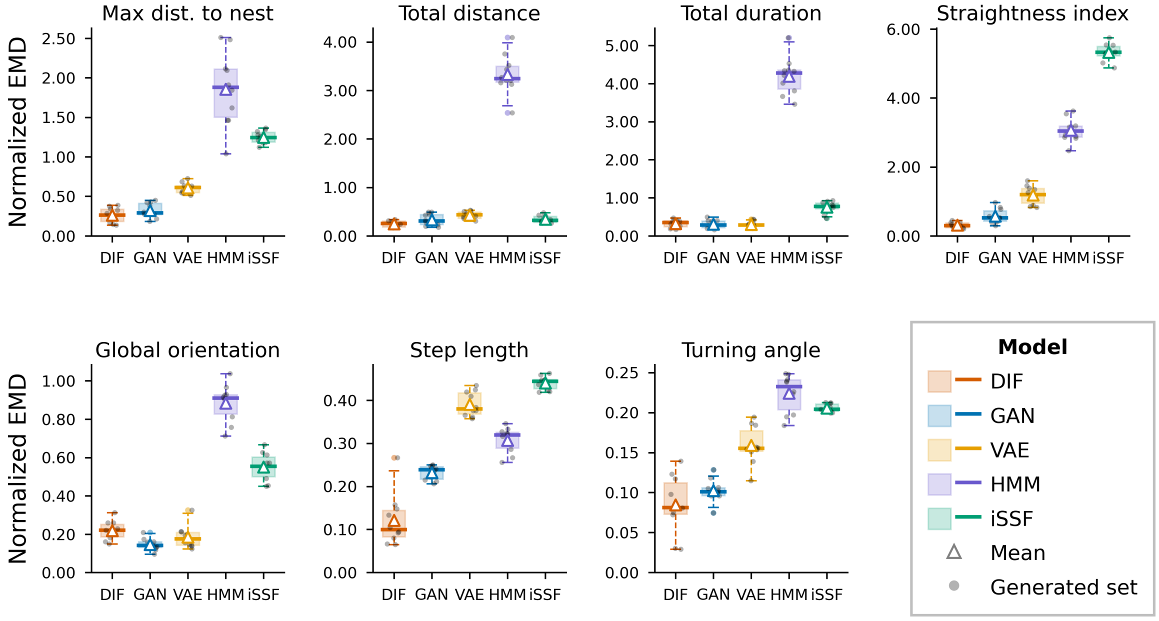


(a) Per-metrics errors.

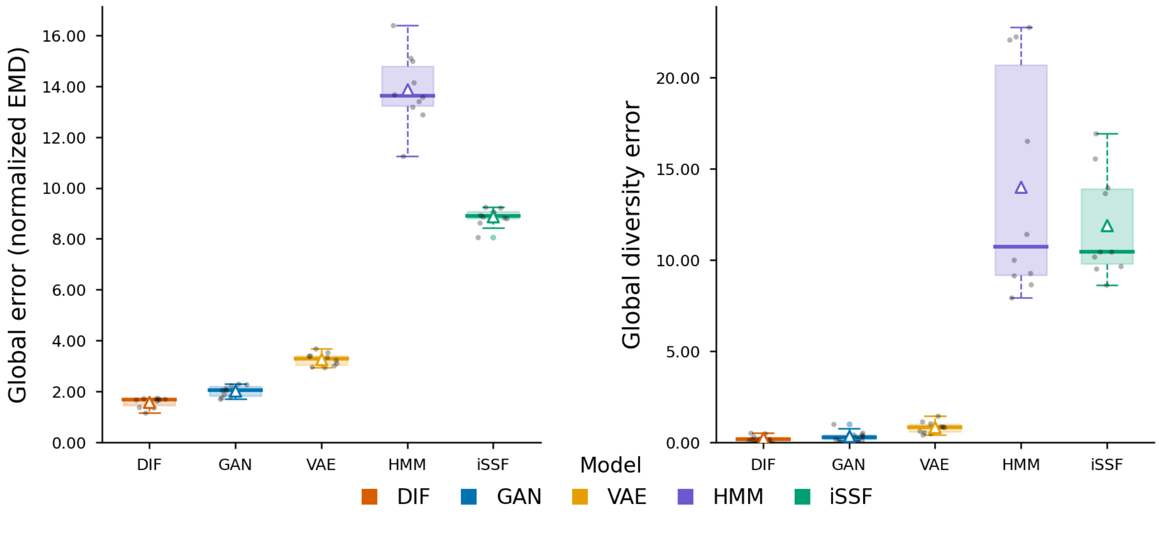


(b) Global scores.

**Fig. 16**: **Model comparison for Peruvian Boobies (*Sula variegata*) at Guañape, Peru ($n$ = 47).** Panels show a) per-metric errors and b) global scores, computed as the Earth Mover's Distance between real and generated distributions across 10 independently generated sets per model.

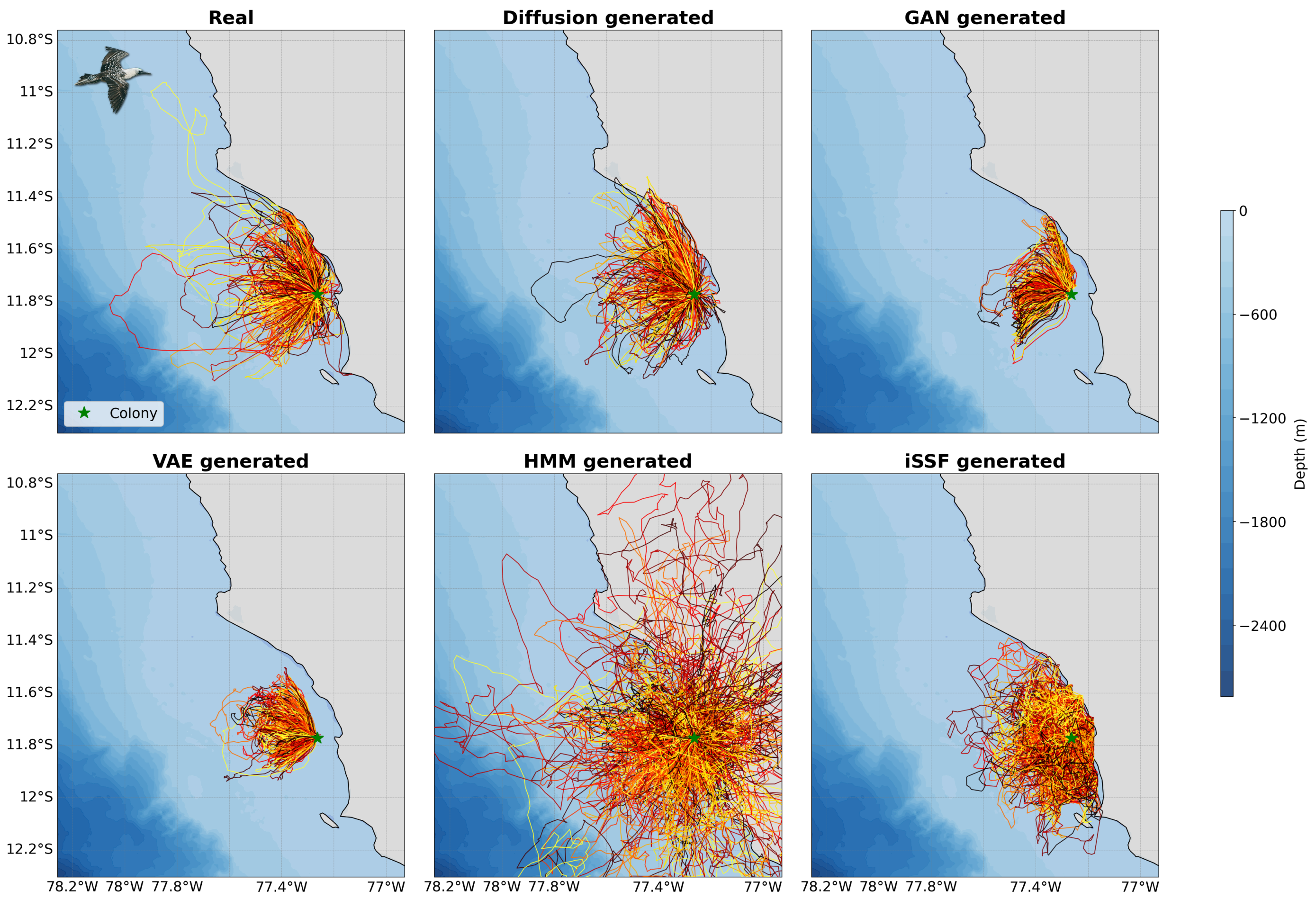


**Fig. 17**: **Real and model-generated trajectories for Peruvian Boobies (*Sula variegata*) at Pescadores Island, Peru.** Each panel shows 412 trajectories. A trajectory is an out-and-back trip from the colony. Colors represent unique trips and are randomized for visual clarity.

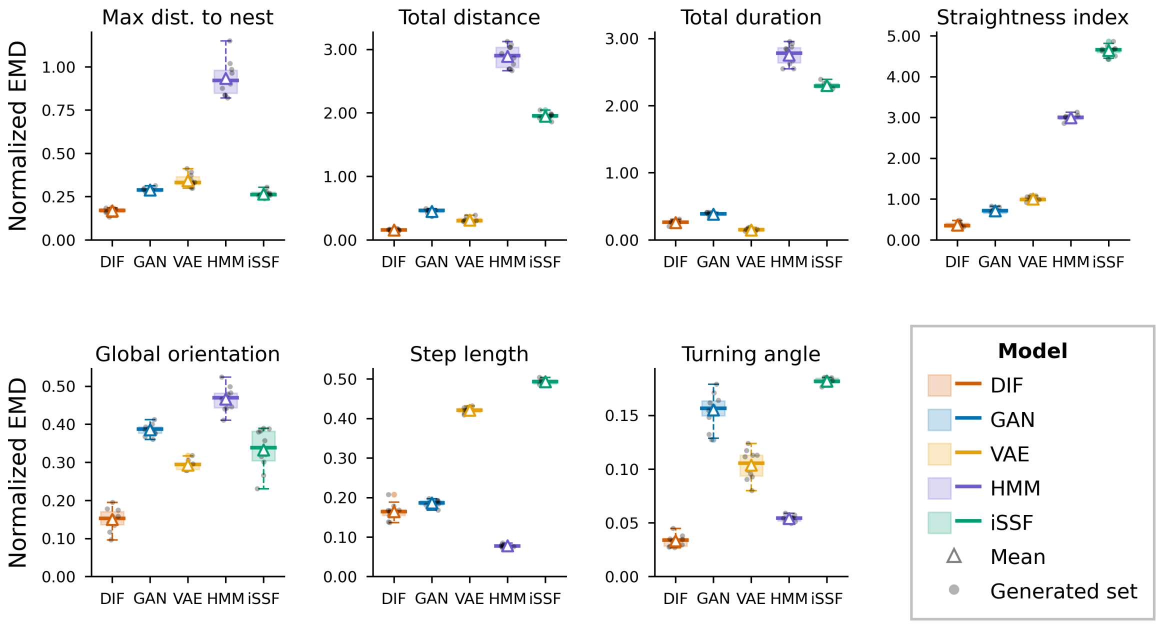


(a) Per-metrics errors.

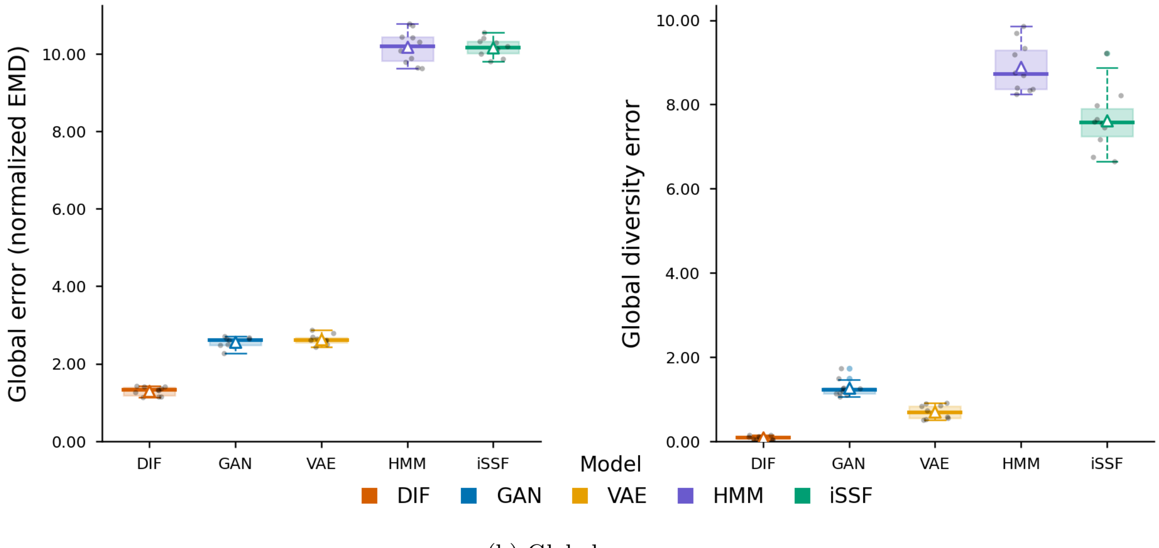


(b) Global scores.

**Fig. 18**: **Model comparison for Peruvian Boobies (*Sula variegata*) at Pescadores Island, Peru ($n$ = 412).** Panels show a) per-metric errors and b) global scores, computed as the Earth Mover's Distance between real and generated distributions across 10 independently generated sets per model.

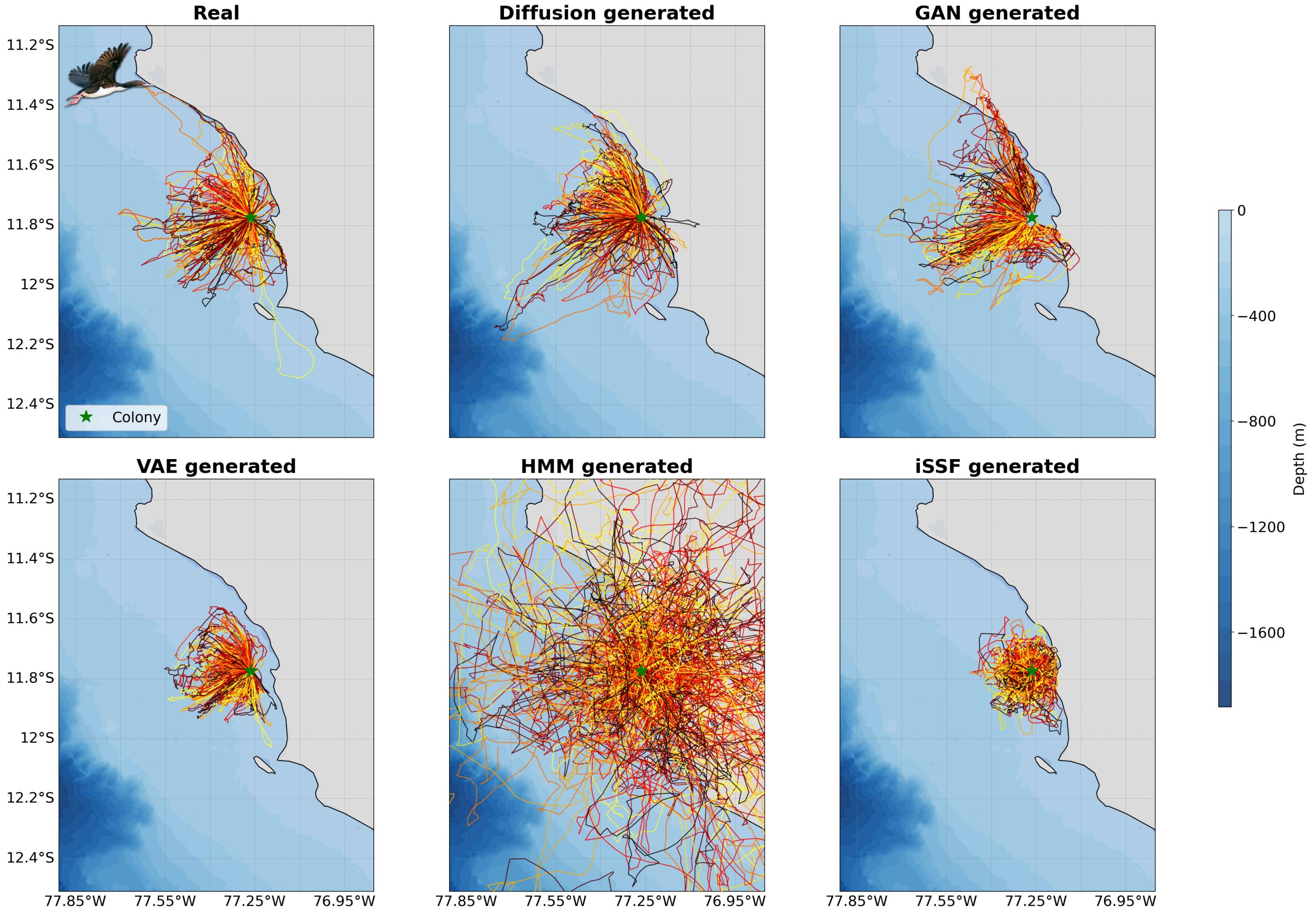


**Fig. 19**: **Real and model-generated trajectories for Guanay Cormorants (*Leucocarbo bougainvillii*) at Pescadores Island, Peru.** Each panel shows 194 trajectories. A trajectory is an out-and-back trip from the colony. Colors represent unique trips and are randomized for visual clarity.

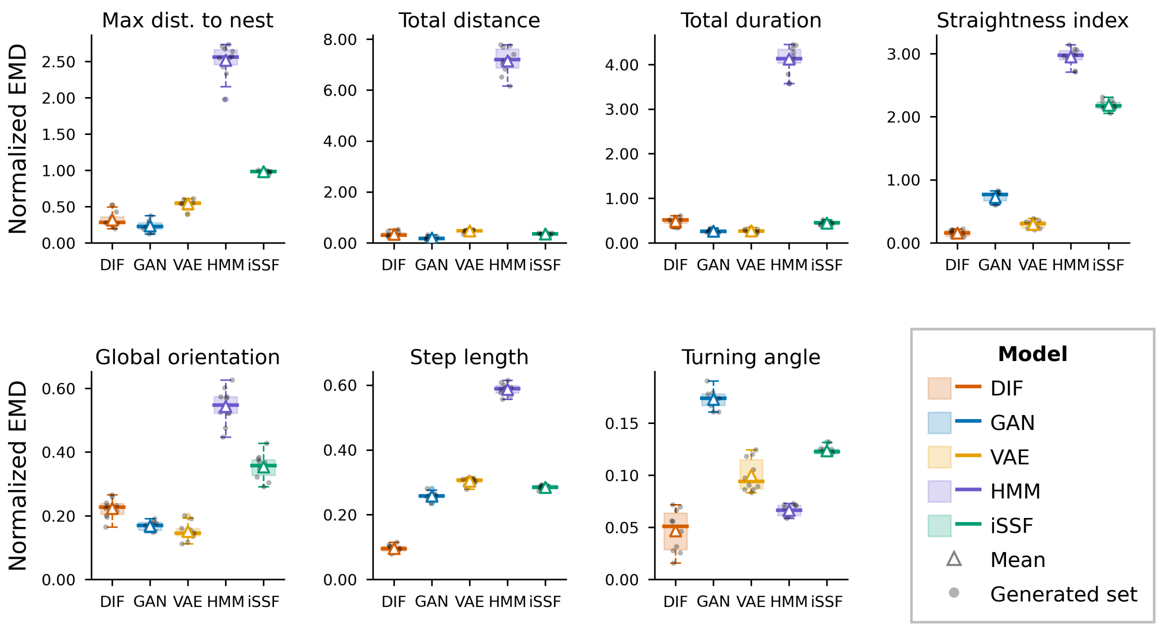


(a) Per-metrics errors.

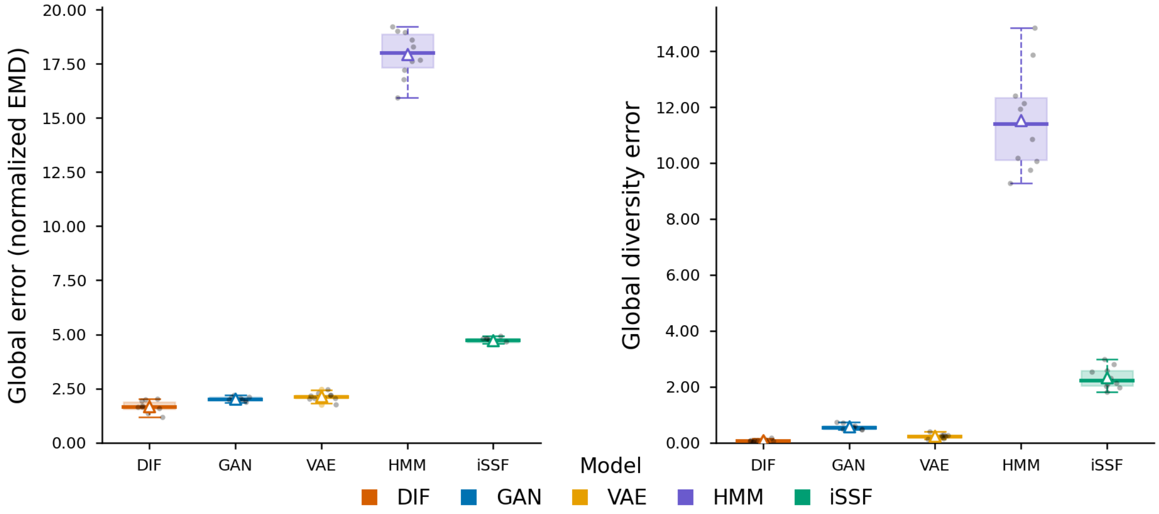


(b) Global scores.

**Fig. 20**: **Model comparison for Guanay Cormorants (*Leucocarbo bougainvillii*) at Pescadores Island, Peru ($n = 194$).** Panels show a) per-metric errors and b) global scores, computed as the Earth Mover's Distance between real and generated distributions across 10 independently generated sets per model.

### C Spatial distribution of HMM inferred states on simulated data

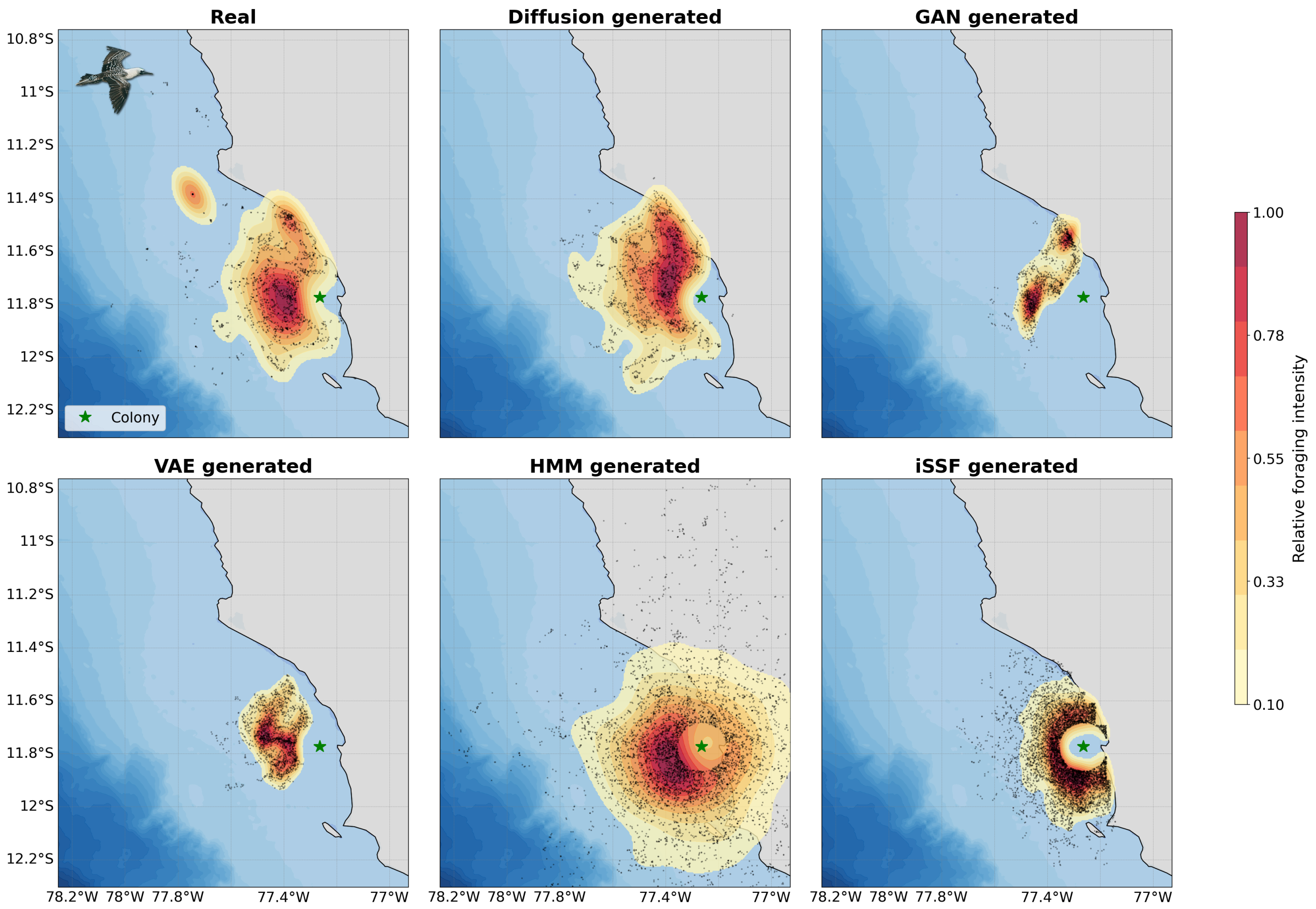


**Fig. 21**: **Spatial distribution of inferred foraging positions for Peruvian Boobies (*Sula variegata*) at Pescadores Island, Peru.** Distributions are shown for real trajectories and those generated by five models: diffusion, VAE, GAN, HMM, and iSSF. Only positions located more than 10 km from the colony are included. Distributions were estimated using Kernel Density Estimation (KDE). Contours represent relative presence probabilities. Black points represent the inferred foraging positions.

### D Behavioral state HMM: emission distributions and state proportions

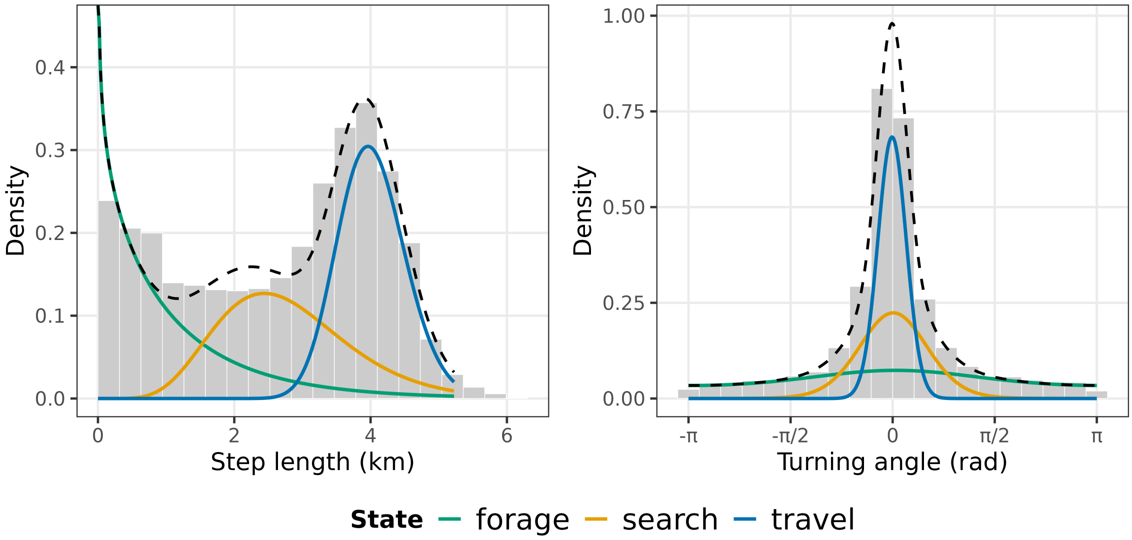


**Fig. 22**: **State-dependent step length and turning angle distributions from the 3-state Hidden Markov Model fitted on real trajectories of Peruvian Boobies (*Sula variegata*) at Pescadores Island, Peru ($n = 412$).** The three states are interpreted as travel (long steps, low turning angle), search (intermediate steps), and forage (short steps, high turning angle). The dotted curve represents the mixture sum across states.

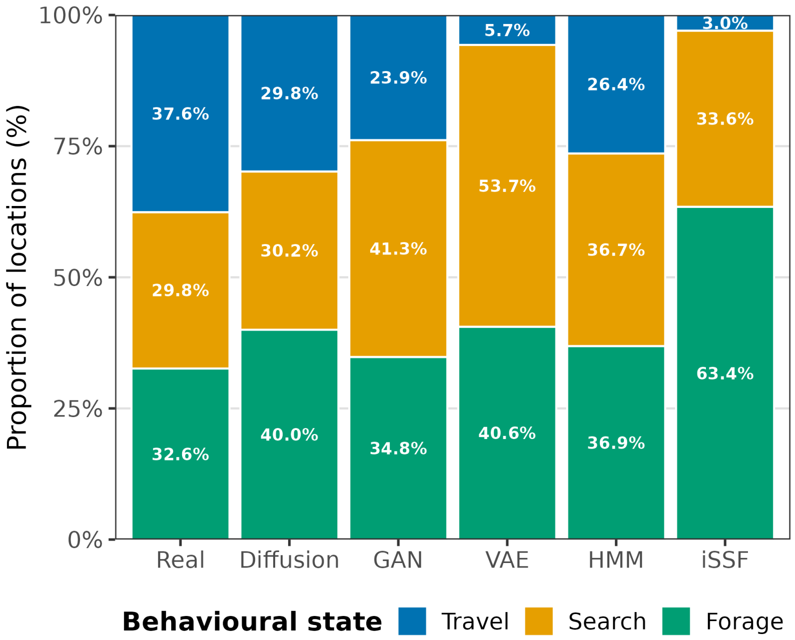


**Fig. 23**: **Proportion of behavioral states across trajectories for real data and five generative models, for Peruvian Boobies (*Sula variegata*) at Pescadores Island, Peru.** States (travel, search, forage) were inferred using the 3-state HMM whose emission distributions are shown in Supplementary Fig. 22.

### E Few-shot adaptation: per-metric error heatmaps

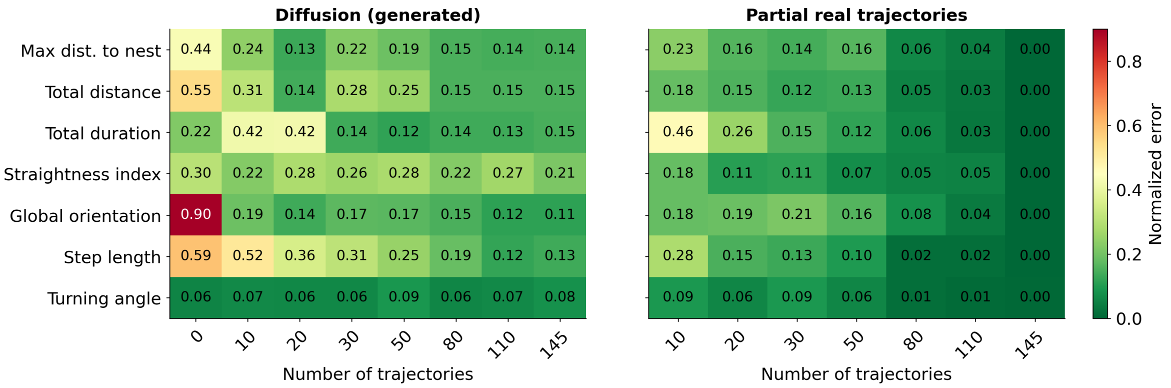


**Fig. 24**: **Per-metric fidelity error heatmaps for the few-shot adaptation of the diffusion model to Brown boobies (*Sula leucogaster*) at Santana Archipelago, Brazil.** Each cell represents the mean error across 10 independent simulation runs, as a function of the number of trajectories used for fine-tuning. This is a heatmap summary of the curves shown in Fig. 7a. Metrics are normalized so that a score of 0 indicates perfect agreement with the real distribution.
